# Mechanistically informed adaptive dosing for cancer immunotherapy using AI-guided decision making

**DOI:** 10.64898/2026.06.09.730783

**Authors:** Ayush Garg, Shyam Sundar Das, Naveen Sivadasan, Arijit Roy, Broto Chakrabarty

## Abstract

Optimizing dose and schedule remains a central challenge in oncology drug development, particularly for immunotherapies where fixed dosing regimens often fail to account for patient specific heterogeneity in tumor–immune dynamics. Here, we present a hybrid quantitative systems pharmacology–reinforcement learning–Monte Carlo Tree Search (QSP–RL–MCTS) framework for personalized immunotherapy dosing that formulates dose selection as a sequential decision-making problem. The approach integrates a mechanistic QSP model of prostate cancer immunotherapy, transcriptomics informed virtual patient populations and data driven AI system comprising reinforcement learning and Monte Carlo tree search. Reinforcement learning is used to learn adaptive generalized dosing policies that optimize treatment outcomes across the population, while Monte Carlo Tree Search provides forward-looking evaluation of RL predicted dosing trajectories to refine patient-specific decisions. On benchmarking against fixed dosing regimens of *ipilimumab*, the remission rate of the proposed model (95.2%) was comparable to the highest fixed dosing regimen of 10 mg/kg per dose while the median total dose (72 mg/kg) of the proposed model designed regimen was comparable to the lowest fixed dosing regimen of 3 mg/kg per dose. The model is generalizable across different dosing protocols and can be extended to predict optimal dose under different therapeutic scenarios. Analysis of the learned dosing trajectories enables stratification of patients into distinct response groups and identifies drug activity rate as the dominant determinant of long-term treatment outcome. These results demonstrate how mechanistically guided artificial intelligence can transform population-level dose optimization into patient-specific, biologically interpretable treatment strategies for precision immuno-oncology.

## Introduction

Dose selection in oncology has historically been guided by the principle that higher drug exposure produces greater antitumor activity until toxicity becomes unacceptable. The current practices of dose optimization in oncology are undergoing a paradigm shift from the traditional maximum tolerated dose (MTD) approaches towards more data-driven, patient-centric strategies that integrate pharmacokinetics (PK), pharmacodynamics (PD), biomarkers, and exposure-response relationships^1,2^. The early stage MTD-based oncology trials used dose-escalation designs (e.g., 3+3 design) to identify the MTD and carry it forward into later trials, assuming a monotonic relationship between dose, efficacy, and toxicity^3–5^. This approach was well suited to cytotoxic chemotherapy, where efficacy and toxicity often increase together over a narrow therapeutic window. With the developments in targeted therapies, monoclonal antibodies and immunotherapy, it has been observed that the maximal biological activity may occur below the MTD and dose–response relationships may plateau before dose-limiting toxicity is observed^6,7^. Unlike chemotherapy induced toxicity, which is typically dose-dependent and acute, immunotherapy is associated with immune-related adverse events (irAEs) that may be delayed, unpredictable, and not strictly correlated with dose, complicating traditional dose-escalation paradigms^8^. In order to address these issues, regulatory authorities have also taken initiatives such as the U.S. FDA’s Project Optimus and the 2024 guidance on oncology dose optimization that encourage randomized dose-ranging studies, model-informed drug development, and comparative evaluation of risk-benefit across doses to ensure that approved doses are not only safe but also optimized for sustained clinical benefit^9,10^.

Personalized dosing in immunotherapy aims to move beyond fixed or weight-based “one-size-fits-all” regimens by tailoring immune checkpoint inhibitor (ICI) exposure to patient- and disease-specific determinants of response and toxicity. Personalized dosing is therefore increasingly recognized as a critical requirement for precision oncology. Inter-patient variability in tumor growth kinetics, immune-cell composition, drug clearance, target sensitivity, and pharmacodynamic response can cause patients receiving the same nominal regimen to experience markedly different treatment outcomes^11,12^. In immuno-oncology, treatment outcome depends not only on drug exposure but also on dynamic interactions among cancer cells, cytotoxic T cells, regulatory T cells, antigen-presenting cells, cytokines, and immune checkpoints within the tumor microenvironment. A uniform dosing regimen may therefore underdose patients with aggressive disease or low drug sensitivity, while overdosing patients who could achieve disease control with substantially lower cumulative exposure^7^. This is particularly important for immune checkpoint blockade, where dose de-escalation or extended-interval strategies have been proposed to preserve efficacy while reducing immune-related adverse events and treatment costs^8,13^.

Quantitative systems pharmacology (QSP) provides a mechanistic framework to address these challenges by integrating pharmacology, disease biology, and systems-level physiology into mathematical models capable of simulating longitudinal treatment response^14,15^. Since its emergence as a model-informed drug development approach, QSP has evolved from pathway-level and pharmacokinetic/pharmacodynamic extensions into multiscale platforms that incorporate molecular signaling, cellular interactions, tissue-level disease processes, and clinical response endpoints^16,17^. Such models have been used to test mechanistic hypotheses, identify predictive biomarkers, compare monotherapy and combination strategies, optimize dose and schedule, and conduct in silico clinical trials using virtual patient cohorts^18–20^.

Model-based simulation can further support personalized dosing by enabling prospective evaluation of alternative dosing regimens in virtual patients before clinical implementation. By initializing QSP models with patient-specific or population-derived biological features, simulations can predict how individual tumor–immune systems evolve under different dose levels and schedules. However, QSP models alone do not automatically solve the sequential decision-making problem of choosing an optimal dose at each treatment time point as the number of possible dosing trajectories increases rapidly with treatment duration, dose granularity, and schedule complexity. Artificial intelligence (AI) provides an efficient complementary approach to QSP models by learning policies for therapeutic actions over the high-dimensional decision space that optimizes the defined objectives through interaction with a QSP model^21–23^. Reinforcement learning (RL) is especially well suited for this adaptive dose optimization because treatment can be formulated as a Markov decision process in which an agent observes the current patient state, selects an action (dosing amount), receives a reward based on tumor control and dose burden, and learns a policy that maximizes clinical objectives^24,25^.

In this study, we developed a hybrid QSP-(RL-MCTS) framework for personalized dose optimization of *ipilimumab* in prostate cancer. The RL agent was trained on virtual population to recommend individualized doses across structured dosing time points, while MCTS was added at inference to prospectively explore future dosing consequences and personalize the RL-derived decisions for an individual patient. The resulting framework was compared against fixed dosing regimens of 3, 5, and 10 mg/kg that have been implemented in clinical trials of *ipilimumab*. The proposed model substantially improved remission and cumulative dose as compared to alternative dosing approaches.

## Results

### Overview of QSP-AI dose optimization workflow

We developed an integrated computational framework that combines a mechanistic quantitative systems pharmacology model of prostate cancer immunotherapy with artificial intelligence–based sequential decision making to enable personalized dose optimization. As illustrated in Fig. 1, the workflow links transcriptomics-informed virtual patients, mechanistic simulation of tumor–immune dynamics, and adaptive decision policies to predict individualized dosing trajectories over the full course of treatment.

**Figure 1.**
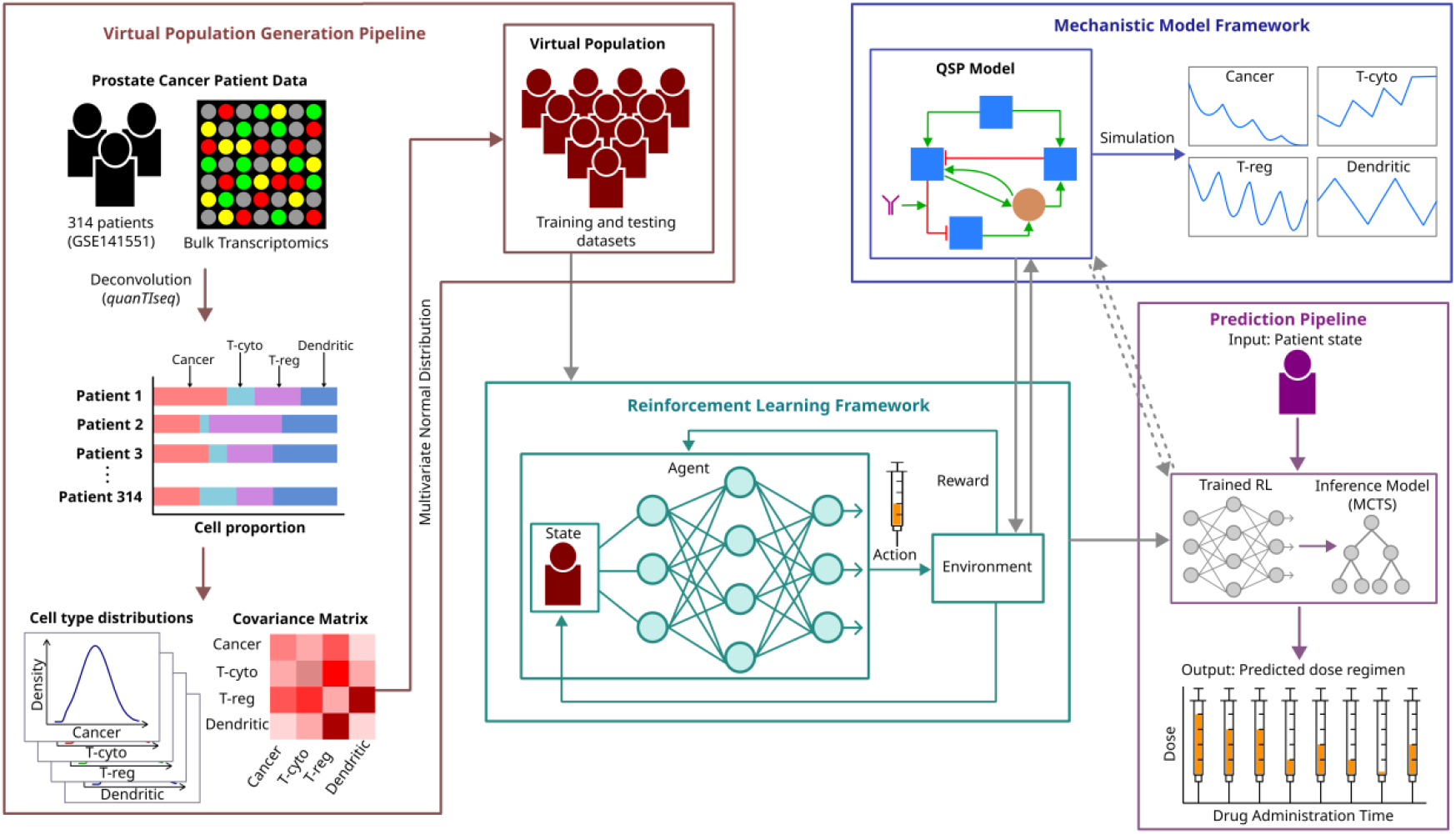
The workflow has four components – virtual population generation pipeline, mechanistic QSP model framework, reinforcement learning framework and the prediction pipeline.

The framework begins with the generation of a heterogeneous virtual patient population derived from bulk tumor transcriptomic data, capturing variability in tumor burden, immune cell composition, and key pharmacodynamic parameters. Each virtual patient is initialized within a previously validated QSP model of androgen-independent prostate cancer treated with the immune checkpoint inhibitor *ipilimumab*, which simulates longitudinal interactions among cancer cells, cytotoxic and regulatory T cells, dendritic cells, cytokines, and drug exposure. This mechanistic model, presented in Fig 2A, serves as a simulator of disease and treatment response under alternative dosing decisions.

**Figure 2.**
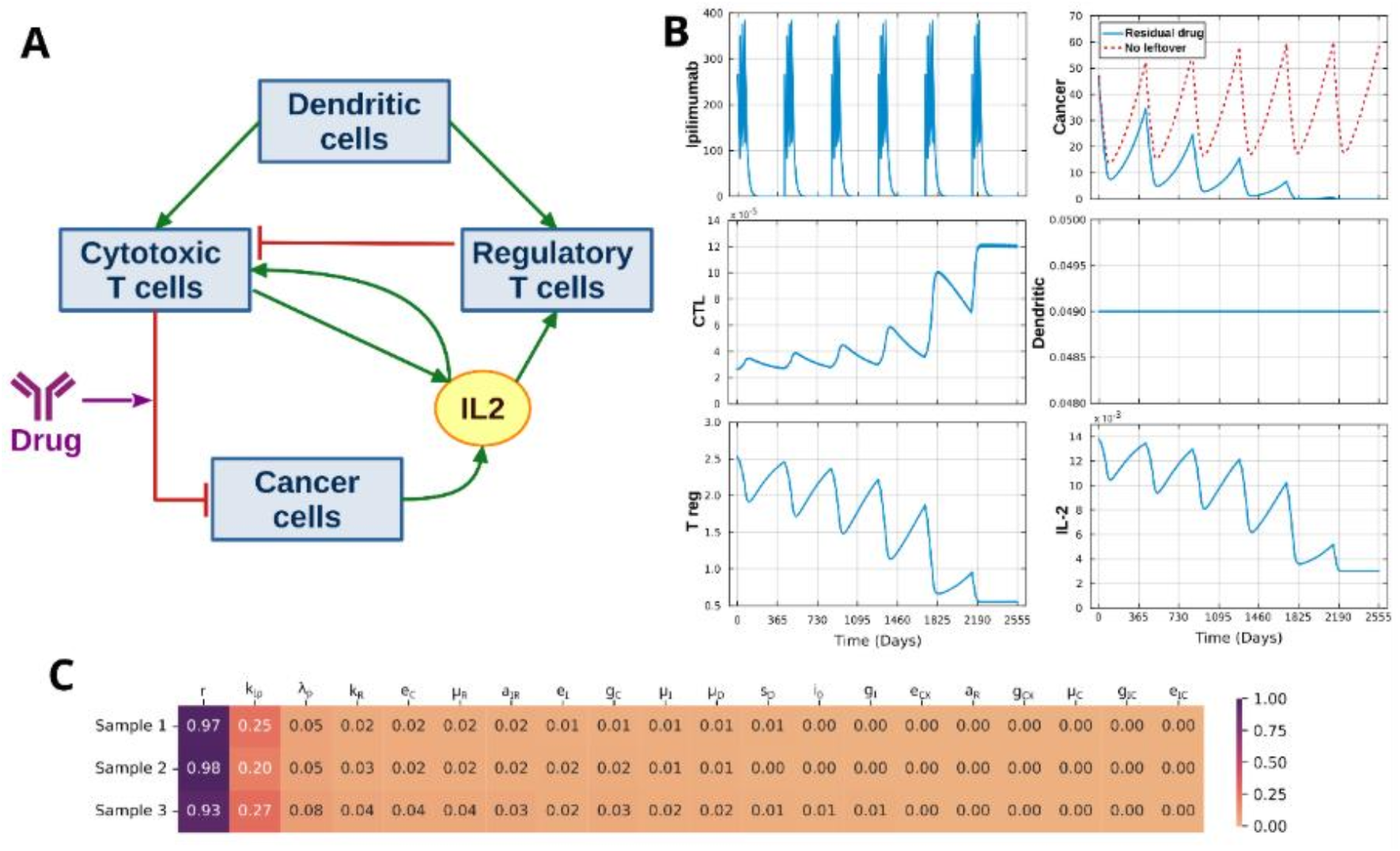
QSP model: (A) QSP model for immunotherapy of prostate cancer by *ipilimumab* (B) Simulation outcome for the duration of 7 years is shown for the drug concentration, cells in the tumor microenvironment – cancer cells, cytotoxic T cells, regulatory T cells and dendritic cells, and the molecular concentration of signalling molecule, IL-2. (C) Global sensitivity values of the top 20 parameters obtained from Sobol method for 3 sample virtual patients are shown.

Personalized dose selection is formulated as a sequential decision-making problem, where reinforcement learning is used to learn a generalized dosing policy across a diverse virtual population. At each dosing time point, the RL agent observes the current patient state as predicted by the QSP model and proposes a dose that aims to maximize long-term tumor control while minimizing cumulative drug exposure. To overcome the inherent population-averaging behavior of generalized policy learning, Monte Carlo tree search is incorporated at inference to prospectively evaluate alternative dosing trajectories and refine decisions in a patient-specific manner.

In the subsequent sections, we first evaluate the performance of the RL agent learned dosing strategies on an independent virtual patient cohort and benchmark them against clinically used fixed dosing regimens. We then analyze how forward-looking planning improves individual treatment outcomes and characterizes the emergent dosing patterns learned by the framework, providing biological and clinical insights into patient-specific treatment strategies.

### Baseline QSP simulations under fixed dosing

The single compartment QSP model for androgen independent prostate cancer treatment with immune checkpoint inhibitor, *ipilimumab*, was adopted from a published human model by Coletti *et al*. ^26^, as shown in Fig. 2A. The mechanistic baseline for treatment response was established by evaluating the behavior of the QSP model under a clinically used fixed-dose regimen of *ipilimumab*. The simulation of the model for an example virtual patient is shown in Fig. 2B. The virtual patient was selected with moderate values of cancer load, proliferation rate and drug response rate. The drug, *ipilimumab*, was administered with a constant dose of 3 mg/kg body weight for 4 times in a year at intervals of 3 weeks for a total simulation time of 7 years^26^. Under this fixed regimen, the cancer cells gradually reduce over the treatment tenure and move towards steady state along with the other cells. The *t*_*1/2*_ for *ipilimumab* is 12.5 days^27^, which means the complete drug clearance from the body takes more than the dosing interval of 3 weeks and the same was observed in the simulation of *ipilimumab* concentration^28^. Some previously published models have also incorporated residual drug in the differential equation governing the drug concentration^29,30^. Therefore, the existing QSP model was modified to incorporate the effect of residual drug in the body at the time of dosing at intervals of 3 weeks. The impact of considering the residual drug can be observed in the cancer progression plot in Fig. 2B, where the progression of cancer cell with and without considering residual drug are shown in blue and red colors respectively.

To characterize the parameters of the QSP model governing the treatment response, global sensitivity analysis of the cancer load was performed by Sobol method to evaluate the importance of the model parameters on the longitudinal simulation of the QSP model^31^. The sensitivity values on three random samples are shown in Fig. 2C. The top three parameters are proliferation rate of cancer (*r*), killing rate of cancer cells by CTL due to *ipilimumab* (*k*_*Ip*_ ) and drug clearance rate (*λ*_*p*_ ). This is consistent with the available knowledge of the behavior of androgen independent prostate cancer and its response against the drug *ipilimumab*. Since the variation in these three parameters have maximal impact on the outcome for patient and the sensitivity of other parameters is negligible, *r, k*_*Ip*_ and *λ*_*p*_ were considered along with the 6 state variables of the QSP model to define inter-patient variability.

While these fixed-dose QSP simulations establish the expected mechanistic behavior of tumor– immune dynamics under standard dosing, they also reveal strong sensitivity of treatment outcome to patient-specific biological parameters, motivating the construction of a virtual patient population to systematically capture inter-individual heterogeneity.

### Virtual population with clinically observed heterogeneity

To train the AI model and evaluate personalized dosing strategies under clinically realistic heterogeneity, we constructed a large virtual patient population that reflects inter-individual variability. Diverse conditions of the tumor microenvironment (TME) and the biological feasible ranges of the parameters were required to capture the variability in cancer progression and treatment response using the QSP model. In recent studies, transcriptomics data analyses have been efficiently used to estimate the TME for immunotherapy QSP models^32–34^. Accordingly, the genome-wide gene expression profiles of primary prostate cancer from a study available at GEO database (accession no: GSE141551)^35,36^ was considered. The relative abundance of the immune cells and cancer cells were estimated using quanTIseq algorithm^37^, which was available at TIMEDB database^38^. The resulting cohort comprised of 314 prostate cancer patients predominantly belonging to Caucasian population (∼92%) and few African Americans (∼8%), covering diverse patient groups in terms of age, Gleason score, PSA and TME, as shown in their respective distributions in Fig. 3A and 3B.

**Figure 3.**
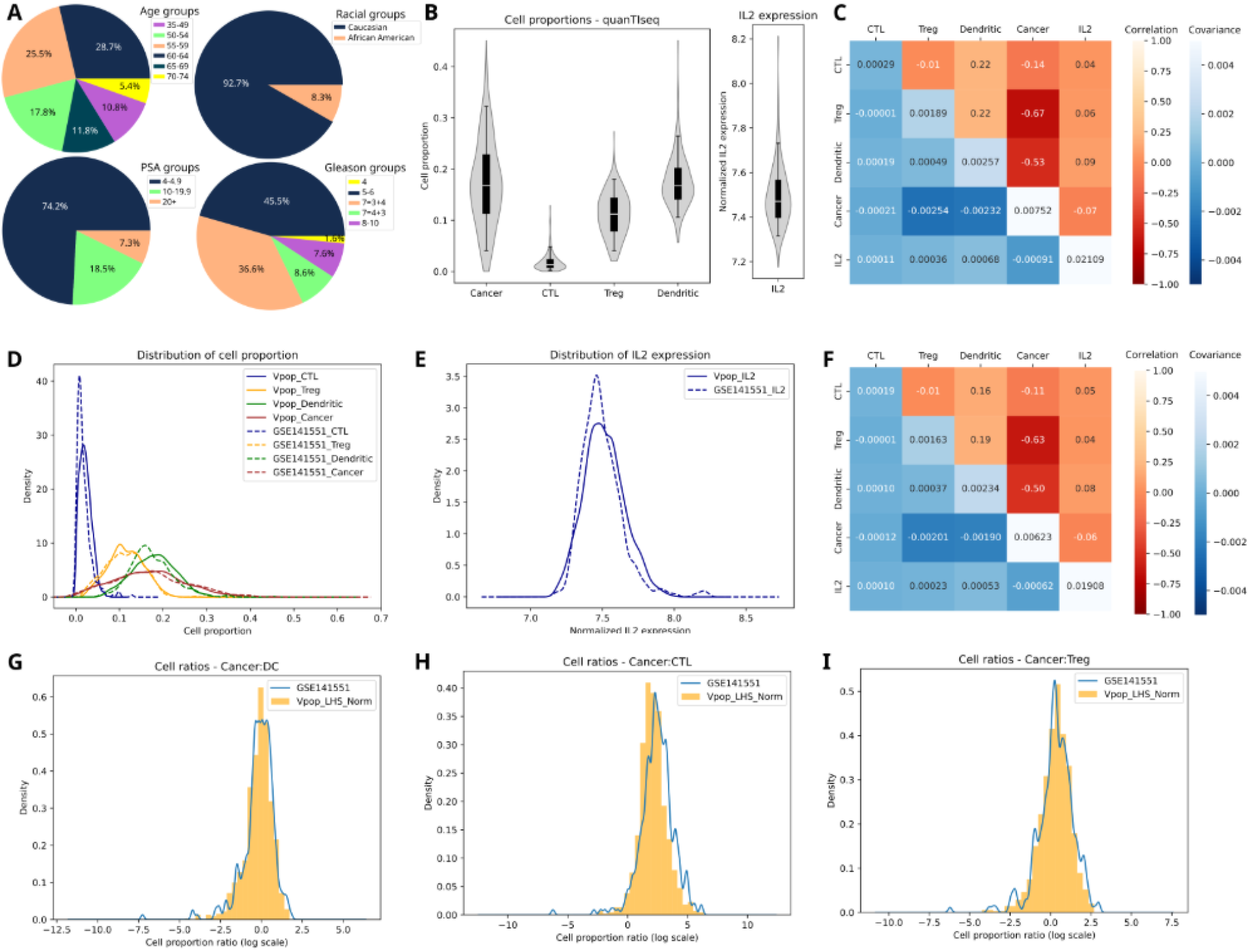
Distribution of features in the reference population of 314 patients from the GEO dataset (GSE141551) are shown in (A)-(C) and that of 10,000 virtual patients are shown in (D)-(I). (A) Pie charts showing distribution across age groups, racial groups, PSA groups and Gleason groups. (B) Distribution of cell proportions and molecular concentration obtained from the gene expression analysis. (C) Correlation and covariance between the features of patients from reference population. (D) Density plots of cell proportions for the virtual population are compared with the reference population. (E) Density plot of concentration of signalling molecule, IL-2, for the virtual population and reference population. (F) Correlation and covariance between the features of patients from virtual population. (G)-(I) show the density plot of ratio of cancer cell proportions to the dendritic cells, cytotoxic T cells and regulatory T cells respectively, for the virtual and reference population.

Considering the GEO dataset as reference, we generated virtual population (VPop) with 10,000 virtual patients following the workflow in Fig. 1. Four approaches for virtual population generation were evaluated -Latin hypercube sampling (LHS), LHS transformed to normal (LHS-N), LHS transformed to truncated multivariate normal (LHS-MN) and simulation-based rejection sampling by Allen et al.^39^ The VPop generated by these methods were assessed against the reference population on two aspects – individual distribution of features which was captured by Kolmogorov-Smirnov (KS) statistic and the relation between the features captured by covariance and correlation matrices. The distribution of cell types and signalling molecule for the reference population are shown in Fig. 3B, and the covariance and correlation matrices are shown in Fig. 3C. The comparative analyses across the methods are provided in the Table T1 and T2 of supplementary file.

Traditionally, LHS is the most popular method for generating virtual population for in-silico clinical trials as it ensures covering the complete parameter space. However, it generates uniform distribution for each feature which neither preserves the distribution of each feature nor the relation between them. This is reflected in the higher KS statistic values and lower covariance and correlation values. In the second approach, the *VPop* was generated by LHS and then transformed into normal distribution keeping the means and standard deviations conserved with that of the reference population. This approach improved the KS statistic for the individual features, but covariance and correlation were not conserved. In the third approach, *VPop* generated by LHS was transformed into truncated multivariate normal distribution keeping the mean, covariance and bounds consistent with the reference population^40^. This method ensured that the individual features followed the unimodal normal distribution like reference population and it was reflected in the low KS statistic values. Additionally, the overall relation between them was also conserved which was reflected in the covariance and correlation matrices. We also generated *VPop* using a simulation-based rejection sampling method. In this method, an initial plausible population was first generated using simulated annealing and *VPop* was selected from it by rejection sampling to match the target population distribution. Although the individual distributions of the features were well-conserved with the reference population, the correlation among features were not preserved. Furthermore, the method exhibited a very low acceptance rate of ∼2% requiring generation of ∼500,000 plausible patients to obtain the desired *VPop* of 10,000 patients. Based on the comparative analyses, the *VPop* generated by LHS-MN was selected for data driven modelling. The distribution of individual features for the virtual population generated by LHS-MN are shown in Fig. 3D-3E, the covariance and correlation in Fig. 3F and the distribution of the ratio of cancer cells to three immune cells – dendritic cells, CTL and regulatory T cells are shown in Fig. 3G-I respectively. It can be observed that the distributions of the virtual population match very well with the reference population. Subsequently, the cell proportions and normalized expression values were transformed into cell counts and molecular concentration by scaling the values to the reference prototype patient given with the QSP model by Coletti et al.^26^

In addition to cellular and molecular concentrations in the TME, the individual variability in tumor progression and treatment response are also governed by the variation in the parameters for individual patients. The sensitivity analysis of the QSP model in Fig. 2C identified proliferation rate of cancer (*r*), killing rate of cancer cells by CTL due to the drug (*k*_*Ip*_ ), and drug clearance rate (*λ*_*p*_ ) as the top three parameters having maximal contribution to the QSP outcome. These three parameters along with the cellular and molecular concentrations were considered to define a virtual patient. Since these parameters are independent of the TME composition, we generated independent normal distributions for these parameters and randomly assigned them to the *VPop* generated by LHS-MN. The bounds of directly measurable parameters, *r* and *λ*_*p*_, were obtained from the literature^26,41^ and the range for *k*_*Ip*_ was determined based on our analysis of the effect of its value on the cancer progression which ensured the coverage of diverse response profiles (Supplementary Figure S2). Finally, a virtual population of 10,000 patients was generated with 8-dimensional virtual patients comprising cancer cells, cytotoxic T cells, regulatory T cells, dendritic cells, IL-2 concentration, *r, λ*_*p*_ and *k*_*Ip*_ . The virtual population was validated by simulating them for 7 years with 3 mg/kg constant dose regimen. The overlapping plot of the simulation in Supplementary Figure S3 confirms that all the variables remain in the physiological bounds throughout the simulation indicating that the population is feasible.

### Reinforcement learning enables adaptive dose optimization but exhibits population-level generalization

The personalized dose optimization for a structured dosing schedule can be conceived as a sequential decision-making problem that follows the properties of a Markov Decision Process (MDP), where personalized doses are predicted for each dosing timepoint of the dosing regimen based on current patient state. Reinforcement learning provides an efficient framework for predicting optimal actions for a MDP and has been efficiently used for the dose optimization of the drug warfarin^25^. A structured dosing schedule for clinical trial of ipilimumab was considered for training the RL model where the 4 doses of the drug is administered at an interval of 3 weeks followed by a gap of 1 year and this dosing pattern is repeated for 6 cycles in 7 years, as shown in Fig. 4A^26^. The objective of our RL model was to learn the optimal dosing policy in the range of 0-10mg/kg for the 24 dosing time points (t1, t2, …, t24) to achieve highest possible patient remission with lowest possible total dose administration. In this study, a patient was considered remitted if the cancer cell count becomes lower than 10-5 billion cells which translates to less than 1 in 10 million prostate cells (as the total prostate volume is of the order of 100 billion cells).

**Figure 4.**
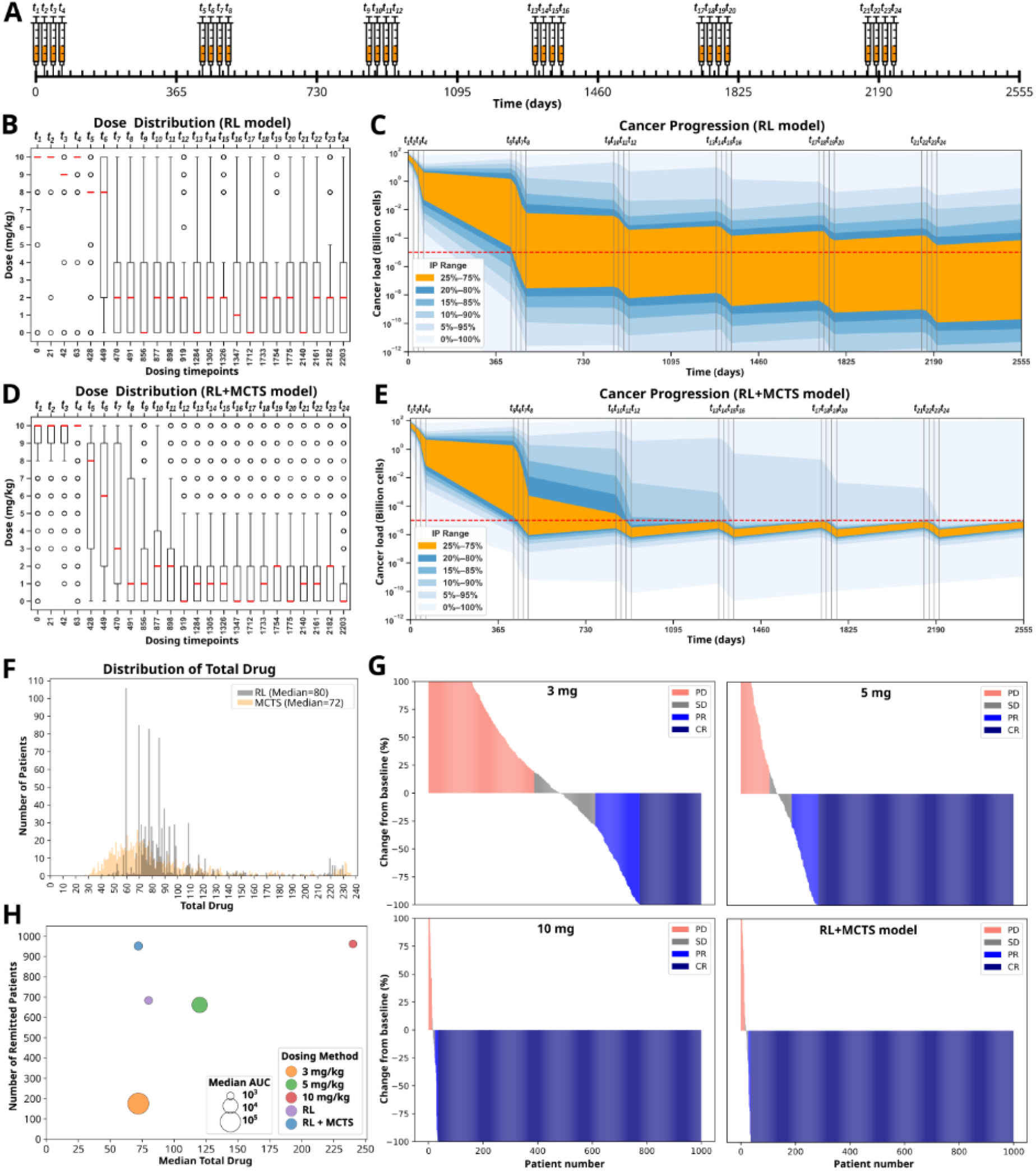
(A) Pictorial representation of the dosing schedule for 7 years – repeating cycle of 4 doses administered at an interval of 3 weeks followed by a gap of one year. (B) Distribution of predicted doses by RL for 24 dosing timepoints. (C) Cancer progression during the 7 years for doses predicted by RL. (D) Distribution of predicted doses by final model (RL+MCTS) for 24 dosing timepoints. (E) Cancer progression during the 7 years for doses predicted by final model. (F) Frequency distribution of total drug predicted by RL and final model. (G) Waterfall plots for the fixed dosing regimens of 3 mg/kg, 5 mg/kg and 10 mg/kg compared with the final model. PD: progressive disease; SD: stable disease; PR: partial response; CR: complete response. (H) Comparison of median total dose and number of patients remitted by the fixed dosing regimens and the final model. All results in (B)-(H) are for virtual population of 1,000 patients (*VPop*_*Test*_ ).

The RL agent was trained using proximal policy optimization (PPO) algorithm on 10,000 virtual patient population (VPop) generated using LHS-MN covering the range of patient parameters and initial states. During training, the RL agent selects a dose for each dosing timepoint form the action space of possible doses and interacts with the QSP model which acts as an environment that simulates the effect of the action on the patient state. The training was performed for ∼750,000 timesteps comprising of ∼31,200 episodes, where the loss and episode reward mean stabilize after ∼500,000 timesteps or ∼21,000 episodes indicating that the convergence of PPO policy learning, as shown in supplementary Fig. S4.

The performance of the trained RL model was evaluated on a set of 1,000 virtual patients (called *VPop*_*Test*_ ) to assess its ability to learn adaptive dosing strategies from mechanistic tumor-immune dynamics. *VPop*_*Test*_ was generated using the same pipeline as *VPop* discussed earlier, ensuring that the patient parameter distributions and covariance matrix remained conserved with the reference population. It was also confirmed that none of the patients in the *VPop*_*Test*_ set is present in the *VPop* set by checking the pairwise Euclidean distance between the virtual patients of both the sets. The distribution of doses predicted by RL for the 24 dosing timepoints on *VPop*_*Test*_ is shown in Fig. 4B and the effect of dose administration on cancer progression during the 7 years for *VPop*_*Test*_ is shown in Fig. 4C. The remission threshold of 10^-5^ billion cells, represented by the red horizontal dashed line, is the optimal target cancer load for our model. The number of patients remitted and the cumulative median dose for test cohort are presented in Table 1. The QSP-RL model predicted dose could remit a reasonable patient count of 684 by administering median cumulative drug of 80 mg/kg. But, the cancer load profiles provided in Fig. 4C, show high diversity indicating the scope of further refinement for most of the patients who were either under-dosed or over-dosed than the optimal dose required to attain the remission threshold. This happened as the RL policy, trained to maximize a terminal reward across a heterogeneous population, favors population-level optimization rather than individualized dose refinement.

**Table 1.**
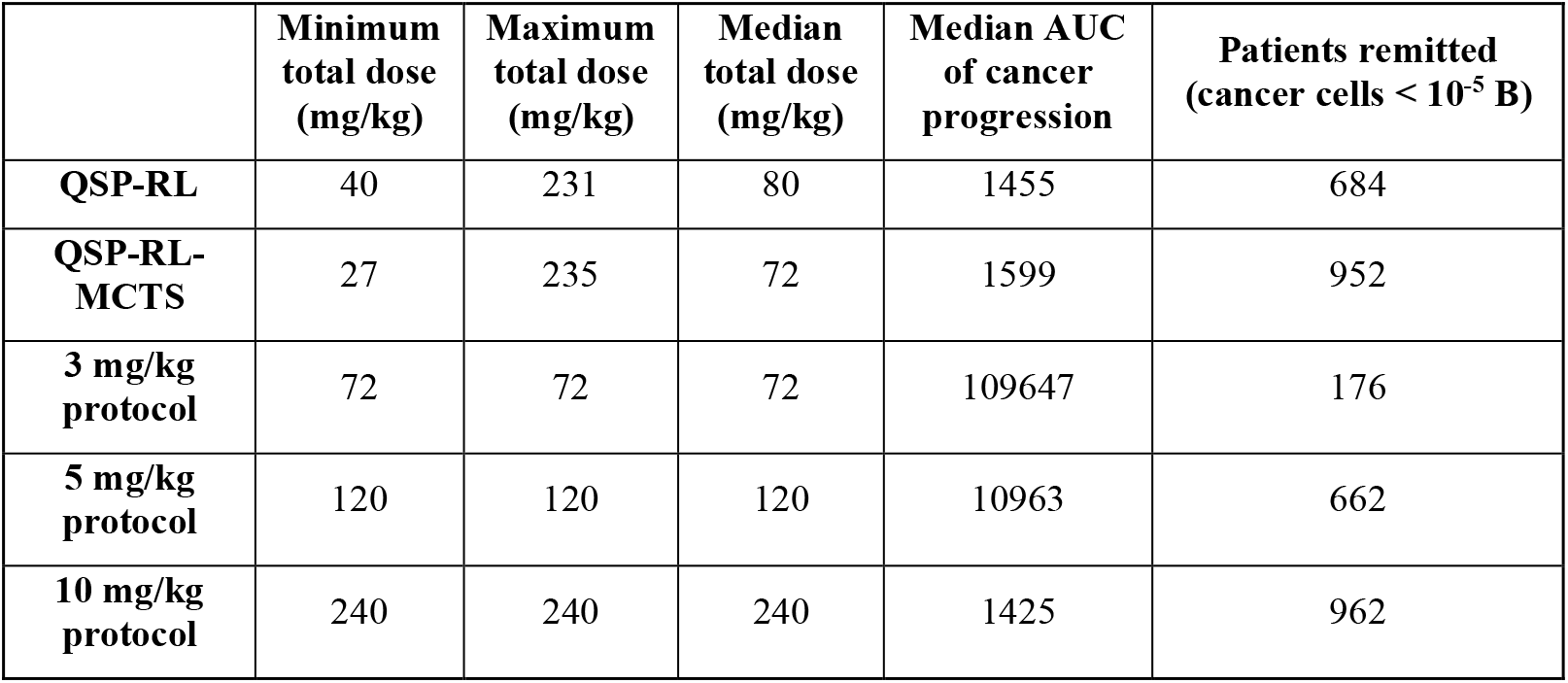
The total dose, AUC (area under the curve) of cancer progression with time and total number of patients remitted by the predictive models (QSP-RL and QSP-RL-MCTS) are compared with the fixed dosing regimens of 3mg, 5mg and 10mg.

| | Minimum total dose (mg/kg) | Maximum total dose (mg/kg) | Median total dose (mg/kg) | Median AUC of cancer progression | Patients remitted (cancer cells $< 10^{-5}$ B) |
| --- | --- | --- | --- | --- | --- |
| **QSP-RL** | 40 | 231 | 80 | 1455 | 684 |
| **QSP-RL-MCTS** | 27 | 235 | 72 | 1599 | 952 |
| **3 mg/kg protocol** | 72 | 72 | 72 | 109647 | 176 |
| **5 mg/kg protocol** | 120 | 120 | 120 | 10963 | 662 |
| **10 mg/kg protocol** | 240 | 240 | 240 | 1425 | 962 |

### Forward-looking Monte Carlo tree search refines RL predictions

A forward-looking Monte Carlo tree search module was integrated during the inference phase of prediction to balance the effect of generalized policy learning of the RL model. While the RL policy provides a generalized mapping from patient state to dose, MCTS refines these recommendations by explicitly simulating and evaluating the downstream consequences of alternative dosing choices over future decision points. Application of the combined QSP-RL-MCTS framework to the independent test cohort resulted in a marked improvement in both treatment efficacy and dosing efficiency.

During the prediction of personalized dosing at each decision time points for a virtual patient, MCTS constructs a decision tree with 5 candidate branches for a depth of 1 year dosing timepoints. Three of these 5 branches were created for 3 most probabilistic doses suggested by the trained PPO policy and 2 additional random doses sampled from the remaining action space. The MCTS evaluates these 5 dosing trajectories and selects the most rewarding action. The dose distribution across the dosing timepoints and the resulting cancer progression for the final QSP-RL-MCTS model on *VPop*_*Test*_ are shown in Fig. 4D and Fig. 4E respectively. The first 4 predicted doses representing the scheduled doses for the first-year, are very high for most of the patients with a median dose of 10 mg/kg, which results in sharp decline in the cancer load for most of the patients. Since the decline depends on the initial state of the patient and parameters such as proliferation rate (*r*) and rate of response to the drug (*k*_*Ip*_ ), highly diverse cancer load profiles are observed across the population at the end of first year. This leads to high variability in predicted dosing profile for the dosing timepoints in the second year. The cancer load stabilizes around the target threshold of 10^-5^ billion cells during the third year, beyond which a maintenance dose is predicted for most of the patients depending on their profile. The frequency distribution of the total drug predicted by QSP-RL model, presented in Fig. 4F, is centred around a limited set of values with very high frequency whereas the frequency distribution is more distributed for the final model. The probable reason for this difference is that the generalized PPO policy learning in RL leads to the same prediction for many virtual patients. In contrast, MCTS refined the RL predictions so that the cancer load converges near the remission threshold for most of the virtual patients, as shown in Fig. 4E. This refinement significantly improved the number of total remitted patients and the cumulative drug exposure, as shown in Table 1. The number of remitted patients increased from 684 in QSP-RL model to 952 in QSP-RL-MCTS model, while the median total drug reduced from 80 mg/kg to 72 mg/kg.

Overall, it was observed that higher doses were preferentially assigned during the early treatment phase to rapidly suppress tumor burden, followed by adaptive dose modulation and maintenance dosing tailored to individual disease trajectories. In contrast to QSP-RL predictions, dose adjustments under QSP-RL-MCTS closely tracked patient-specific tumor dynamics, preventing both under-treatment of resistant cases and over-exposure in patients achieving early disease control. Together, these results demonstrate that forward-looking planning resolves key limitations of generalized policy learning by enabling individualized exploitation of mechanistic tumor–immune simulations. The integration of MCTS with RL transforms population-level dose optimization into a patient-specific decision-making process, yielding superior efficacy–exposure tradeoffs relative to RL alone and providing the basis for personalized immunotherapy dosing.

### Comparison of personalized and fixed dosing regimens

We next benchmarked the personalized QSP-RL-MCTS dosing strategy against clinically used fixed-dose regimens of *ipilimumab* (3, 5, and 10 mg/kg) to assess its ability to balance treatment efficacy and drug exposure. Fixed regimens were simulated over the same treatment horizon for the independent virtual patient cohort with 1000 virtual patients, and outcomes were evaluated using identical remission criteria. As expected, increasing fixed dose was associated with higher remission rates but also substantially greater cumulative drug exposure (Table 1). The QSP-RL-MCTS model predicts a vast spectrum of personalized cumulative drug ranging from 27 mg/kg to 235 mg/kg, depending on the patient profile. On one hand it prevented over-dosing in patients that could achieve remission with much lower dose and at the same time it ensured higher doses for the patients with aggressive disease profile. The waterfall plots in Fig. 4G show that the response profile of the final model is substantially improved compared to the 3 mg/kg and 5 mg/kg dosing regimen and is similar to that of 10 mg/kg dosing regimen. Further, Fig. 4H shows that the predicted personalized dosing regimen could achieve a response profile comparable to that of the highest fixed dosing regimen of 10 mg/kg while administering a median total dose of 72 mg/kg which is equal to the total administered dose of 3 mg/kg dosing regimen.

We further analysed the patients remitted by the three fixed dosing regimens and compared their outcomes under the QSP-RL and final QSP-RL-MCTS models. Table 2 shows that 176 patients were remitted under 3 mg/kg dosing regimen, out of which 173 were remitted by the QSP-RL predicted dose while3 patients did not achieve the remission threshold. Out of the 173 patients remitted by QSP-RL model, the total administered dose for 145 patients was within the total dose of 3 mg/kg regimen, i.e. 72 mg/kg, while for the remaining 28 patients the QSP-RL predicted total dose was more than 72 mg/kg. It can be argued that these 28 patients could have been remitted with dose within 72 mg/kg since they were remitted by the 3 mg/kg fixed dose regimen. Also, for the 3 patients that were not remitted, the total predicted dose was less than 72 mg/kg, suggesting that they could have been administered more drug to remit. On the other hand, all 176 patients were remitted by the final model with a total dose less than 72 mg/kg. This demonstrates that MCTS significantly improves the outcome in both the ways – higher patient remission and lower total dose. Similar trends were observed for the patients remitted by 5 mg/kg and 10 mg/kg regimen, where all the patients remitted by the final model were predicted with total dose less than the respective fixed dosing regimen. It should be noted that while considering the patients for 5 mg/kg and 10 mg/kg, the patients that were remitted by the lower dose regimen were not considered, i.e. 176 patients that were remitted by 3 mg/kg regimen were not considered in the 5 mg/kg analysis and 662 patients were not considered for the analysis of 10 mg/kg analysis. Among the 300 patients remitted by the 10 mg/kg fixed dosing regimen, only 10 patients were not remitted by the final model. The cancer progression and predicted dosing for these patients are shown in supplementary figure Fig. S5. These patients have high cancer progression rate and lower drug response rates, though 5 of these patients attained a near-threshold remission level (<10^-4^ billion cells).

**Table 2.**
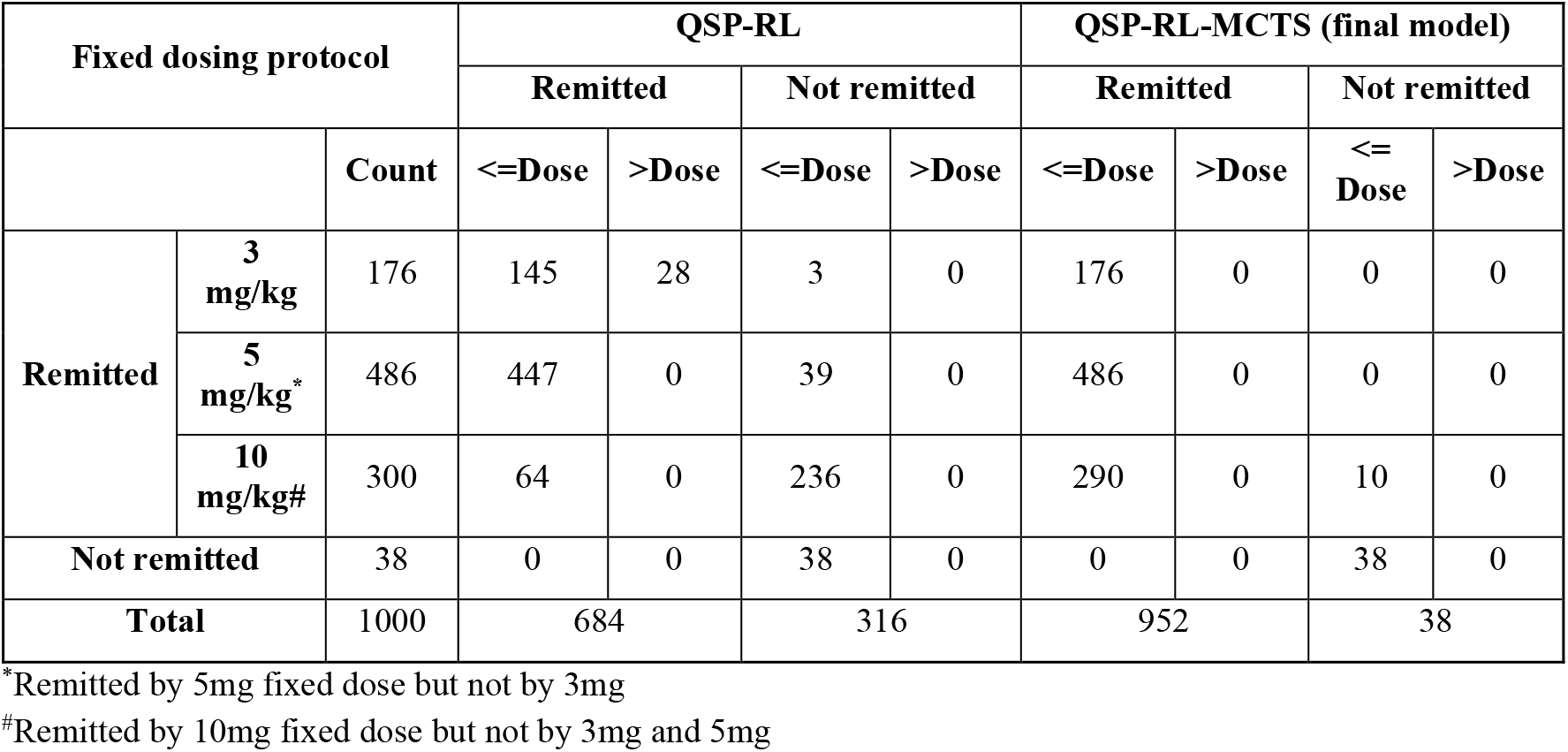
The patients remitted by the fixed dosing regimens of 3 mg/kg, 5 mg/kg and 10 mg/kg are compared with the outcomes of the two hybrid models in terms of number of remitted patients and total dose.

### Learned dosing trajectories stratify patients into distinct treatment response groups

To further interpret the personalized dosing strategies learned by the QSP-RL-MCTS framework, we analyzed whether the predicted dosing trajectories encode coherent patterns that stratify patients into clinically meaningful groups. The total drug predicted by the final model range between 27 mg/kg to 235 mg/kg, reflecting the different strata of patients in the set. We clustered the 1,000 patients based on the dosing profile instead of the general practice of clustering the population based on patient features and associating the patient features to the response profiles. The predicted dose for 24 timepoints was represented as a 24-dimensional vector for each patient and those dosing profiles were clustered by applying *k*-means clustering (*k* = 4) based on the Euclidean distance between the predicted dose profiles. This alternate approach of grouping the patients based on the predicted dosing regimens implicitly considers the patient features that are important in the context of the therapeutic response. The distribution of patients among the 4 clusters is shown in Fig. 5A. Each cluster exhibits a distinct dosing pattern that can be clearly observed from the dose distribution of the 24 timepoints for the 4 clusters in Fig. 5B. Although the RL model was trained only to optimize a terminal reward without any constraints towards individual dosing patterns across the dosing timepoints, it is interesting to observe that dosing patterns for three out of four clusters can be distinctly divided into aggressive and maintenance dosing phases, which can be clearly observed in the median doses for each timepoints in Fig. 5D.

**Figure 5.**
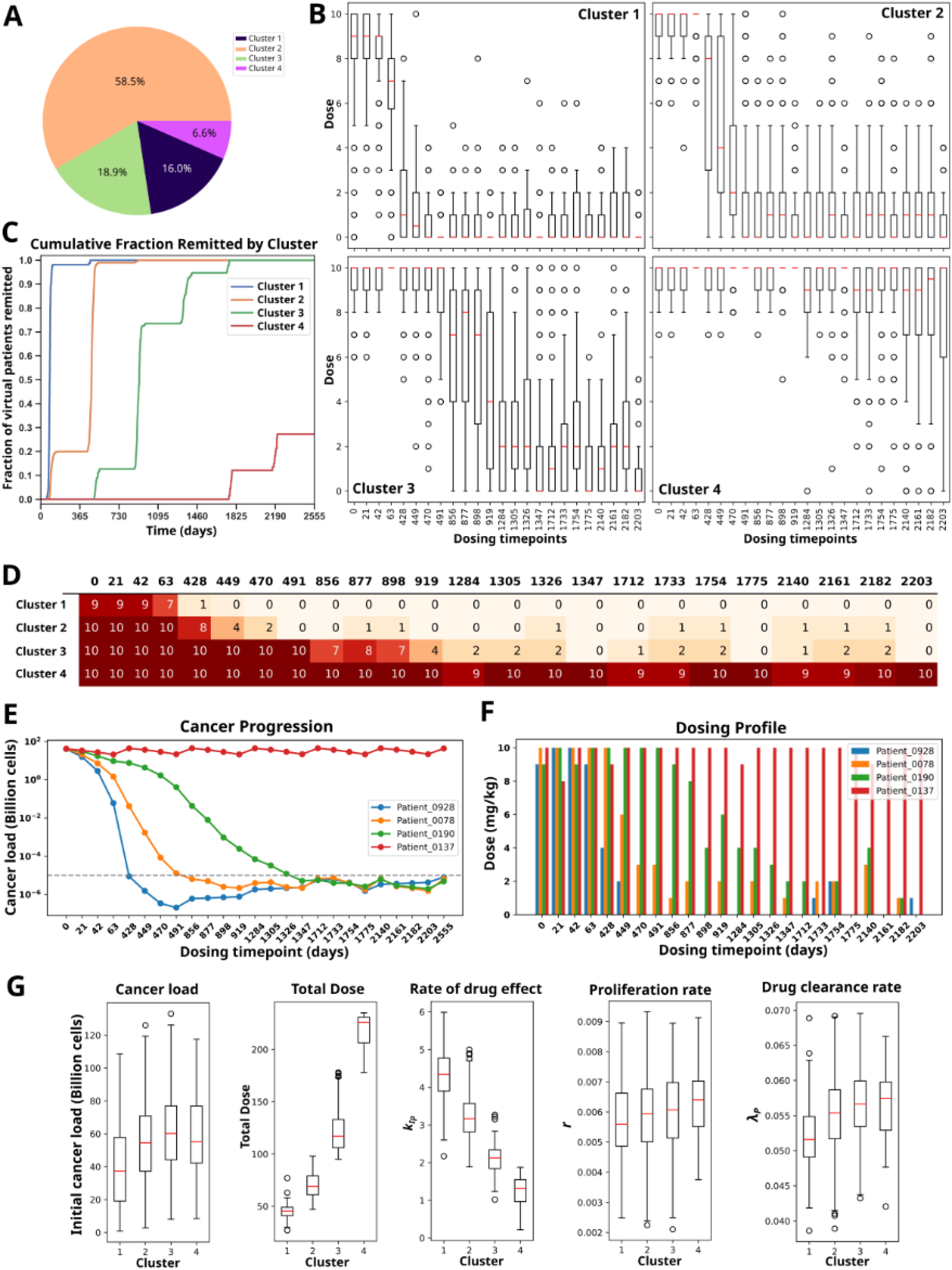
Cluster analysis based on dosing profile. (A) Membership distribution of *VPop*_*Test*_ in the 4 clusters. (B) Dose distribution in the 24 dosing timepoints. (C) Cumulative fraction of patients remitted for each cluster over time. (D) Median dose for the 24 dosing timepoints for the clusters. (E) Cancer load trajectory for 4 example virtual patients belonging to the 4 clusters with similar baseline cancer load. (F) Dosing profile for the 24 dosing timepoints for the example virtual patients in E. (G) Boxplot showing distribution of cancer load, total dose, *k*_*Ip*_, *r* and *λ*_*p*_ for the clusters.

Analysis of remission dynamics revealed substantial differences in treatment timelines among clusters (Fig. 5C). For the Cluster 1, doses for *t*_*1*_ to *t*_*3*_ belonging to the first year are aggressive, it starts reducing from *t*_*4*_ and becomes significantly low in the maintenance phase from dosing timepoints *t*_*5*_ onwards. The pattern of dosing changes in Cluster 2, where *t*_*1*_ to *t*_*4*_ are very high aggressive doses with less variation, dose starts reducing from *t*_*5*_ to *t*_*7*_ with high variability among patients and remains low in maintenance phase beyond *t*_*8*_ . The aggressive dosing is predicted for *t*_*1*_ to *t*_*8*_ in Cluster 3, followed by one year of highly variable dose in third year from *t*_*9*_ to *t*_*12*_ leading to a maintenance phase of low dosing from *t*_*13*_ onwards. Cluster 4 remains in aggressive dosing for all the timepoints with no maintenance dosing.

The long-term effect of the treatment is well captured in the prolonged simulation time-period of 7 years. Fig. 5C shows the remission rate of the clusters with time, which can guide in deciding the duration of treatment required for different strata of patients. The patients of Cluster 1 require treatment for 1 year followed by very minimal maintenance dose at distant time intervals whereas the patients of Cluster 2 require higher doses for longer time-period of around two years beyond which a low maintenance is sufficient to maintain remission. The patients of Cluster 3 attain remission after 5 years which includes aggressive dosing for 2 years, intermediate dosing for one year and remaining period of maintenance dose. Among the virtual patients of Cluster 4, only ∼27% of patients attain remission in the entire duration of 7 years. However, the long simulation shows that the remission in the aggressive disease cases only start towards the end of 5 years after a prolonged high dose of *ipilimumab*.

It is intuitively obvious from the above analysis that the virtual patients in the Clusters 1 to 4 are in the increasing order of difficulty of treatment using the drug, *ipilimumab*. To evaluate the role of initial cancer load on the dose prediction and treatment outcomes, we considered four virtual patients with similar initial cancer load but belonging to the four different clusters. The simulation of the cancer load starting from the same initial load of ∼40 billion cells is shown in Fig. 5E and the predicted dose for the 24 timepoints for these patients are shown in Fig. 5F. The dosing pattern for all the four cases are consistent with the median dose of the respective cluster shown in Fig. 5D and the cancer progression is also distinctively different in the four cases irrespective of the initial cancer load. The distribution of initial cancer load in Fig. 5G also shows that the range of initial cancer across the four clusters are similar.

These results demonstrate that personalized dosing trajectories learned by the framework naturally stratify patients into distinct response groups with interpretable treatment requirements and remission timelines. This dosing-trajectory–based stratification provides a clinically relevant understanding heterogeneity in immunotherapy response and forms the basis for mechanistic interpretation of treatment difficulty in subsequent analyses.

### Features responsible for cluster differentiability

We examined the baseline tumor burden and key QSP model parameters to identify the biological factors underlying the distinct dosing profile–based patient clusters described earlier. For this purpose, the three parameters that were part of the virtual population - proliferation rate of cancer (*r*), killing rate of cancer cells by CTL due to ipilimumab (*k*_*Ip*_ ) and drug clearance rate (*λ*_*p*_ ) were evaluated. The distribution of *r, k*_*Ip*_ and *λ*_*p*_ are shown in Fig. 5G. It can be observed that the distribution of total drug in the patients was inversely correlated to *k*_*Ip*_, both of which have distinct pattern for the four clusters while the other factors such as *r* and *λ*_*p*_ have similar distribution in all the clusters. These observations indicate that the dosing pattern is primarily governed by the *k*_*Ip*_ which is a composite parameter determined by the several factors such as effect site concentration, absorption rate, binding affinity and activity.

### Generalizability of the AI model

To demonstrate the generalizability of the proposed approach, the pre-trained RL model was applied to an alternative structured dosing schedule used in clinical trials of the drug *ipilimumab*^*42,43*^. In the alternative schedule, shown in Fig. 6A, the drug is administered for 4 times at an interval of 3 weeks followed by maintenance dosing every 3 months. The predicted dosing regimen on *VPop*_*Test*_ resulted in remission of 972 patients. The boxplot of the dose distribution across 30 dosing timepoints and the corresponding cancer load progression for *VPop*_*Test*_ dataset are presented in Fig. 6B and 6C respectively. High doses with median dose of 9 mg/kg are administered to most of the patients in first four dosing points which results in immediate decrease in the cancer load. The variability of dose across the papulation is highest for the next 4-5 dosing timepoints where doses were predicted based on the patient profiles. By the end of the second year, cancer load for most of the patients becomes close to the remission threshold which leads to prediction of maintenance dose for the remaining timepoints.

**Figure 6.**
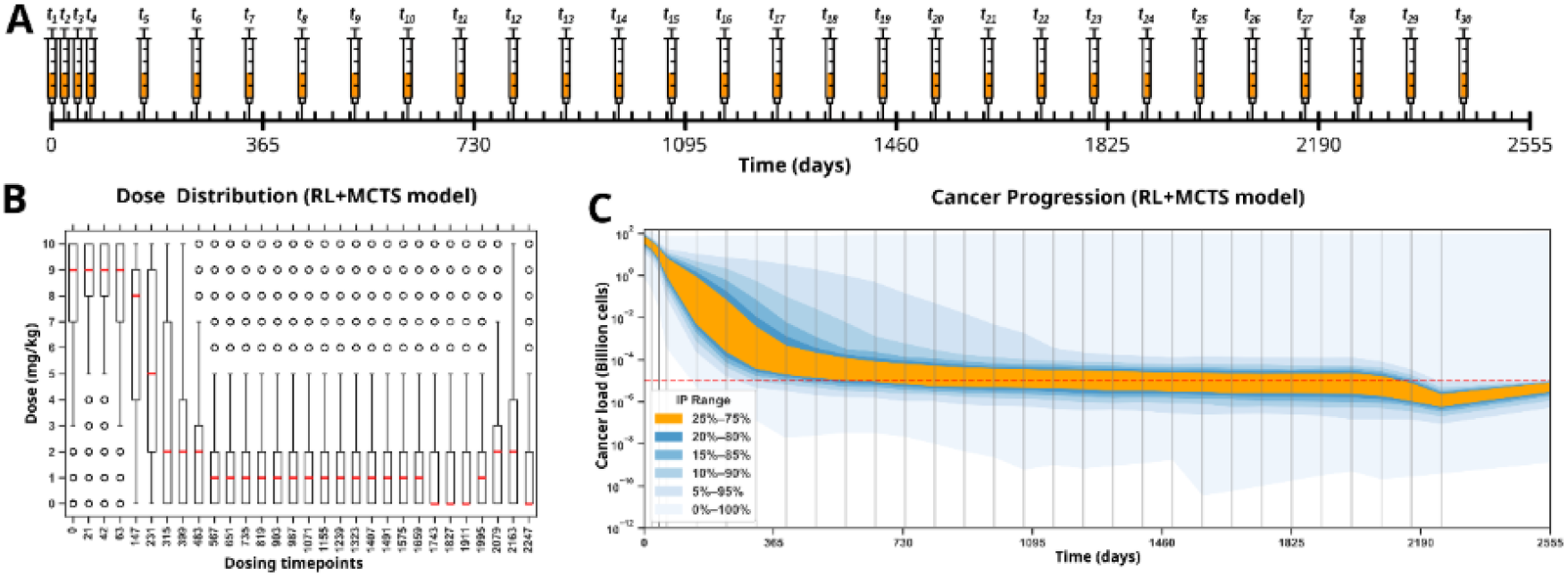
(A) Pictorial representation of the dosing schedule for 7 years – 4 doses administered at an interval of 3 weeks followed by dose administration at every 3 months. (B) Distribution of predicted doses by RL for 30 dosing timepoints. (C) Cancer progression during the 7 years for doses predicted by RL.

## Discussion

In this study, we have demonstrated a proof of concept of integrating mechanistic QSP model with data-driven AI models to build a framework for predicting personalized optimal dose. We integrated a QSP model for the immunotherapy of prostate cancer through immune checkpoint inhibitor drug, *ipilimumab*, within the environment of the reinforcement learning framework. The RL model demonstrated reasonable performance on the test cohort, achieving remission in ∼68.4% of virtual patients. However, since the RL was trained using generalized PPO policy, the dose predictions for the initial timepoints were conservative with very low variation. Except very few exceptions, the predicted doses for the timepoints *t*_*1*_ to *t*_*5*_ for all the patients in Fig. 4B were same. Also, similar cumulative doses were predicted for different group of patients which is reflected in the high frequency for few drug regimens in Fig. 4F. This resulted in high variation in the outcome of cancer load progression shown in Fig. 4C indicating suboptimal personalization. To address these limitations, a forward-looking MCTS layer was integrated on top of the RL policy. The MCTS explored the future outcome trajectories to select the most rewarding dose, which led to higher variation in the dose predictions for individual timepoints (Fig. 4D), lower variation in the cancer load progression (Fig. 4E) and a distributed frequency of total predicted drug (Fig. 4F). Consequently, the remission rate improved significantly to 95.2% with much lower median total dose of 72 mg/kg. The remission rate of patients under the final model, QSP-RL-MCTS, is comparable to the maximal fixed dosing regimen of 10 mg/kg where 96.2% patients were remitted after administering substantially higher total drug of 240 mg/kg to all the patients. In contrast, the median total drug predicted by the final model is comparable to the minimal fixed dosing regimen of 3 mg/kg where administration of a total drug of 72 mg/kg results in only 17.6% remission. Also, for all the 952 patients that were remitted by the final model, the total predicted dose was lower than the fixed dosing regimen required to remit them, as shown in Table 2, highlighting the efficiency of the personalized strategy.

A limitation of the final hybrid model is that the predicted doses remained conservative for ∼5% of the virtual patients in the test set, which were not remitted. The total predicted doses for these 48 virtual patients were lower than the maximum possible total dose of 240 mg/kg. However, 10 of these virtual patients were remitted by the 10 mg/kg fixed dosing schedule and the remaining 38 patients could have attained improved outcomes close to the remission threshold on administration of highest possible dose. Under the final model predicted dosing, the cancer load for 4 out of these 10 patients goes marginally below the remission threshold after the final dose but increases back again in the final year (supplementary Fig. S5), while 6 others have higher final cancer load. For these difficult-to-treat patients, the final model does not predict the highest possible total drug as it compensates the marginal reward gain for improvement in cancer load with the negative reward for higher drug. This trade-off results in underestimation of the optimal dose in a small subset of patients, indicating an area for potential refinement of the reward design and long-term outcome weighting.

The virtual population of 1,000 virtual patients in the test set were generated based on a reference population of 314 real patients with clinically localized prostate cancer. As shown in Fig. 3A, ∼83.8% patients belonged to Gleason grade group 1 and 2 with low grade cancer, ∼8.6% belonged to Gleason grade 3 with intermediate growth and ∼7.6% constituted highly aggressive cancer form. Also, prostatectomy was the primary treatment for all the patients and the samples for gene expression were extracted from the prostatectomy. Therefore, the analysis of the outcomes from this study provides a basis to understand and design the treatment protocol for patients with the clinically localized prostate cancer post prostatectomy.

Together, this analysis demonstrates that the proposed dose optimization framework not only personalizes dosing decisions but also reveals biologically meaningful strata of treatment difficulty driven by pharmacodynamic responsiveness. By translating latent biological variability into distinct, interpretable dosing strategies, the framework provides a mechanistic bridge between patient heterogeneity and adaptive treatment planning, completing the link from personalized decision making to underlying disease biology.

The proposed integrated framework provides a proof-of-concept for integrating mechanistic disease models, omics-informed virtual populations, reinforcement learning, and forward-looking Monte Carlo tree search to predict optimal dose for immunotherapy. The workflow can be extended to other therapeutic areas with different disease biology and treatment mechanisms.

## Methods

### Prostate cancer QSP model

The QSP model for androgen independent (also called castration resistant) prostate cancer treatment with immune checkpoint inhibitor, *ipilimumab*, was adopted from a published human model by Coletti *et al*.^26^ It is a single compartment model that captures the cellular dynamics of the tumor microenvironment and its regulation by the cytokine signalling molecule interleukin-2 (IL-2), as shown in Fig. 2A. The model focused on the androgen independent prostate cancer dynamics post administration of the first line of treatment, i.e. androgen deprivation therapy. Therefore, the system was considered devoid of androgen signalling and androgen dependent prostate cancer cells. The model consists of 6 state variables which include the drug concentration, IL-2 concentration and count of 4 cell types – cancer cells, cytotoxic T cells, regulatory T cells and dendritic cells. The longitudinal evolution of these state variables was modelled using 6 ordinary differential equations (ODE) that are governed by 21 parameters. The ordinary differential equations (ODEs) for these 6 variables are given in Table 3 and the details of 21 parameters are given in Table 4.

**Table 3.**
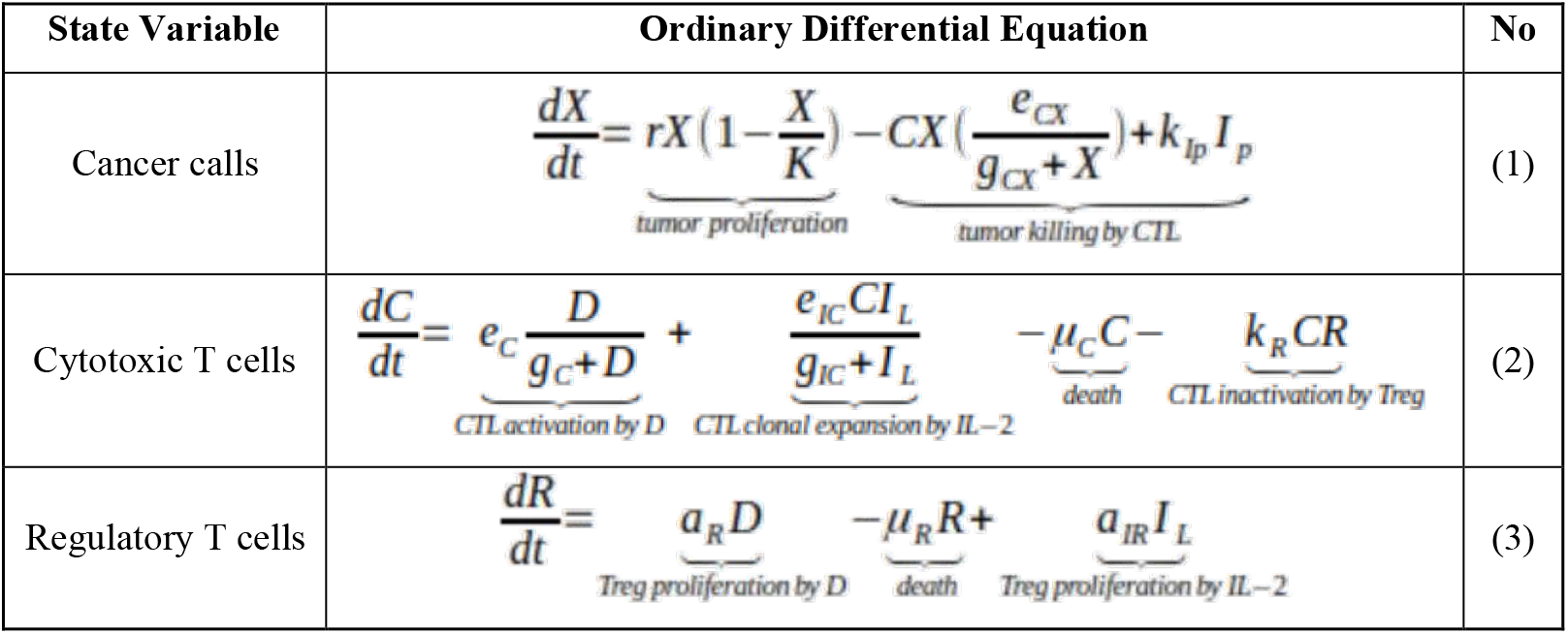
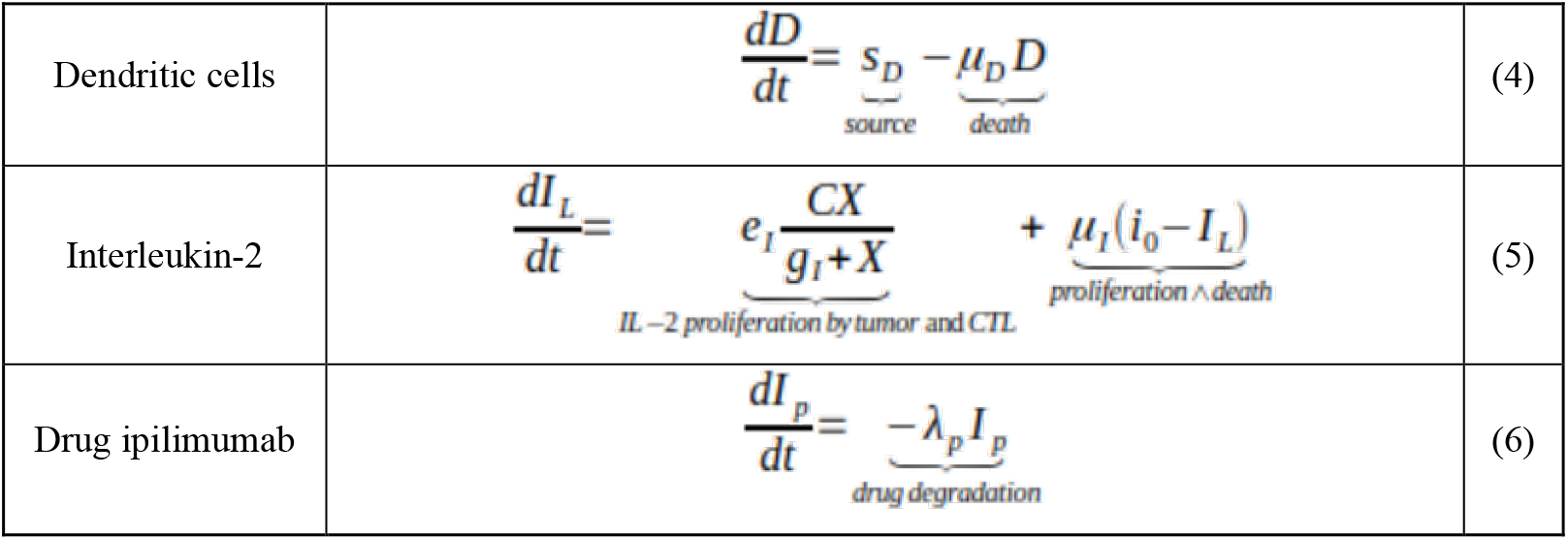
Ordinary differential equations of the QSP model.

**Table 4.** The parameters of the QSP model.

| Parameter | Description | Value | Source |
| --- | --- | --- | --- |
| $r$ | Proliferation rate of cancer cells | 0.006 (± 66.6%) day<sup>-1</sup> | 41 |
| $k_{Ip}$ | Killing rate of cancer cells by CTL due to ipilimumab | 0 - 6<br>(day * mg * billion cells)<sup>-1</sup> | [Figure S2] |
| $\lambda_P$ | Drug clearance rate from the body | 0.055 (± 50%) day<sup>-1</sup> | 26 |
| $K$ | Maximum cancer carrying capacity | 107 billion of cells | 26 |
| $e_{CX}$ | Killing rate of cancer cells by CTL without drug | 0.75 day<sup>-1</sup> | 41 |
| $e_C$ | Activation rate of CTL by Dendritic cells | 0.02<br>*billion of cells/day* | 41 |
| $e_{IC}$ | Activation rate of CTL by IL-2 | 0.1245 day<sup>-1</sup> | 41 |
| $e_I$ | Activation rate of IL-2 by CTL | 5000<br>ng/ (ml * day * billion of cells) | 41 |
| $k_R$ | Inactivation rate of CTL by Tregs | 32.81 (day * billion of cells)<sup>-1</sup> | 26 |
| $a_R$ | Activation rate of Treg by mature Dendritic cells | 0.072 day<sup>-1</sup> | 45 |
| $a_{IR}$ | Activation rate of Treg by IL-2 | 131.26 (ml * billion cells) /<br>(ng * day) | 26 |
| $i_0$ | Baseline level of IL-2 | 0.00299 ng/ml | 26 |
| $s_D$ | Source of dendritic cells | 0.00686 billion cells/day | 26 |
| $g_{CX}$ | CTL saturation level for cancer cells inhibition | 10 billion cells | 41 |
| $g_C$ | Dendritic cell saturation level for T cell clonal expansion | 0.4 billion cells | 41 |
| $g_I$ | Cancer cell saturation level for CTL stimulation | 10 billion cells | 41 |
| $g_{IC}$ | IL-2 saturation level for T cell clonal expansion | 1000 ng/ml | 41 |
| $\mu_C$ | CTL death rate | 0.03 day<sup>-1</sup> | 41 |
| $\mu_R$ | Treg death rate | 0.72 day<sup>-1</sup> | 45 |
| $\mu_D$ | Dendritic cell death rate | 0.14 day<sup>-1</sup> | 41 |
| $\mu_I$ | IL-2 clearance rate | 10 day<sup>-1</sup> | 41 |

### Sensitivity Analysis of QSP model

Sobol method was used to perform global sensitivity analysis of the QSP model^31,44^. The cancer load which is the primary output of the QSP model, was considered as the output variable for sensitivity analysis. The parameter representing the cancer carrying capacity (*K*) determines the upper steady state of the QSP model and the system reaches steady state when cancer load reaches *K*. Therefore, *K* was not considered for sensitivity analysis and was assigned a fixed value of 150. A range has been defined for the remaining 20 parameters for executing Sobol. The range for the parameter *r* was obtained from the literature, for *k*_*Ip*_ it was obtained from the experiments given in supplementary figure Fig. S2 and the range for the remaining 18 other parameters were obtained by considering a variation of ±30% of their values in the Table 4. We used three sets of initial values for the state variables for more reliable analysis.

### Virtual population pipeline

#### Gene expression analysis

The gene expression data from Fred Hutchinson (FH) Cancer Research Center PrCa study was obtained from Gene Expression Omnibus database (accession number: GSE141551)^35,36,46^. It is composed of European-American male residents of King County, Washington, who were diagnosed with prostate cancer either in 1993-1996 or in 2002-2005^47,48^. These patients were diagnosed with clinically localized adenocarcinoma of the prostate and underwent radical prostatectomy as primary therapy. Specimens were extracted from the prostatectomy to obtain expression from the Human HT-12 v4 BeadChip (Illumina) platform. The raw read counts were transformed into reads per kilobase of transcript per million reads mapped (RPKM) and normalized using quantile normalization.

The bulk RNA-seq data was used to estimate the relative abundance of cancer and immune cell proportions using quanTIseq algorithm^37^. It is a deconvolution algorithm that uses constrained least squares regression to estimate the proportions of the immune cells in the tumor microenvironment from the RNA-seq data. The quanTIseq estimates were obtained from TIMEDB database^38^. From the 503 samples reported in the GEO study GSE141551, 314 were filtered by discarding the samples with missing PSA values, PSA less than 4 (indicating healthy individuals), estimated cytotoxic T-cell proportions and cancer cell proportions close to zero,

### Virtual population generation methods

#### Latin hypercube sampling (LHS)

LHS is a stratified sampling technique designed to efficiently explore high-dimensional spaces by enforcing uniform coverage of each variable’s range. In this method, the range of each variable is divided into equiprobable intervals and one sample is drawn from each interval. Then, the samples across dimensions are paired through random permutations, which ensures uniform distribution for each variable^49,50^.

#### LHS transformed to normal (LHS-N)

The distribution of the variables of the reference population were unimodal normal distributions, as shown in Fig. 3D and Fig. 3E. Therefore, to preserve the properties of each individual variable, in this method, the samples generated by LHS were transformed into normal distribution for each dimension by conserving the mean and standard deviation of the reference population. The generation of samples by LHS and then transformation into normal distribution led to preservation of ranking in each dimension, which ensured maximal diversity in the generated virtual population.

#### LHS transformed to truncated multivariate normal (LHS-MN)

In the third method, samples were first generated by LHS and then transformed into truncated multivariate normal distribution keeping the mean and covariance consistent with the reference population, which ensured the relation between the variables were captured in the generated population. The algorithm considers the bounds of the variables from reference population to generate the truncated multivariate normal distribution^40^. This helped in handling skewed distributions and prevented generation of negative values.

#### Simulation-based rejection sampling

Simulation and rejection sampling based method by Allen *et al*.^*39*^ was used as the fourth method to generate virtual population. At the first step, multi-dimensional samples were generated by random sampling followed by optimization of these samples using simulated annealing such that the outcome of the QSP simulation of the samples remained within physiological range of the state variables. These biologically viable samples were called plausible patients. Rejection sampling was then applied on the plausible population to select the final virtual population that follows the properties of the reference population.

### Evaluation of virtual population generation methods

The virtual populations were evaluated on similarity of the individual features and overall population distribution against the target real population of prostate cancer patients.

#### Kolmogorov-Smirnov statistic

The KS statistic is a goodness of fit measure that quantifies the largest vertical distance between two cumulative distribution functions (CDFs). It ranges between 0 to 1, where lower values indicate higher similarity between the two distributions.

#### Covariance matrix

It captures the direction of linear relationship between all the pair of variables, showing how they change together. Positive values indicate the two variables move in the same direction whereas negative value indicates the two values move in opposite directions.

#### Correlation matrix

Pearson’s correlation coefficient measures the strength and direction of a linear relationship between two variables, ranging from -1 to 1. It is obtained by normalizing the covariance of two variables by the product of their standard deviations.

### Reinforcement learning

Reinforcement learning is a branch of machine learning which focuses on determining optimal solution in a sequential decision-making task formulated as Markov Decision Process (MDP). The mathematical framework for the RL consists of decision point, state, action, environment, reward function and an algorithm to find optimal solution in the form of optimal policy^51^. The RL algorithm employs an agent to learn optimal sequence of actions through interactions with an environment and iteratively optimises a policy to maximize the cumulative reward received while exploring the possible action space^52,53^. The QSP model, is considered as the environment in this study, which receives the current state of the patient and the action taken by the RL agent and performs the simulation to generate the next state^51^.

#### Decision point (*t*)

In the context of the dose optimization, decision points represent the time points at which the drug was administered. Multiple sets of decision points including those derived from clinical trial protocols of ipilimumab^*42,43,54,55*^ and proposed by Coletti *et al*.^26^ were evaluated. In the clinical trial protocol, the decision points are day 1, 22, 43 and 64, followed by every 12 weeks. While the schedule proposed by Coletti *et al*. follows the same four initial dosing (day 1, 22, 43 and 64) with this initial cycle is repeated after 365 days for five additional times.

#### State (S)

The virtual patients are represented by the six state variables and three selected parameters of the QSP model. Apart from these state variables and parameters, the state of the RL comprises a history of actions and corresponding cancer load and normalized time progression. The state at n-th decision point is represented by

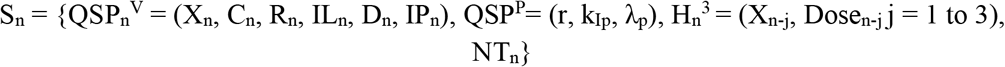

where QSP_n_ ^V^ represents the value of the QSP variables, cancer cell count (*X*), cytotoxic T cell count (*C*), regulatory T-cell count (*R*), IL-2 concentration (*IL*), dendritic cell count (*D*) and drug concentration (*IP*), at *n*^*th*^ decision points. QSP^P^ represents the time independent values of the three selected QSP parameters (r: proliferation rate of cancer, k_Ip_ : killing rate of cancer cells by CTL due to the drug, and λ_p_ : drug clearance rate) considered in the virtual patient. H_n_ ^3^ is the history which contains last three dose and cancer cell counts. *NT*_*n*_ is the normalized time corresponding to *n*^*th*^ decision point.

#### Action (*a*)

The action or decision is referred to the amount of *ipilimumab* to be administered to a patient at a decision point. The minimum possible drug amount considered for administration is 0 mg/kg and the maximum amount considered is 10mg/kg. A discreate action space is considered in this study and for *n*^*th*^ decision point it is defined as

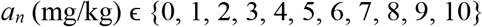

#### Reward function (*R*)

The reward function computes a scalar reward value which indicates the extent of loss or gain for the action *a*_*n*_ at *n*^*th*^ decision point in the state *S*_*n*_ . The primary aim in this study is to cure the prostate cancer patient i.e. reduce the cancer load to <10^-5^ billion cells and desirable aim is to administer as less cumulative drug as possible considering the high cost and toxicity of the immunotherapy drugs. Terminal reward, which is computed at the end of an episode/trajectory is considered to be better suited for this dose optimization study. Accordingly, the terminal reward function (equation (1)) was defined in terms of two quantities, *reward*_*cancer*_ representing the reward corresponding to the cancer load at the end of an episode and *reward*_*dose*_ representing the cumulative administered dose for a virtual patient.

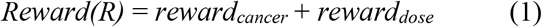

if (*X*_*t=2555*_ > 10^-5^):

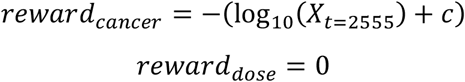

else:

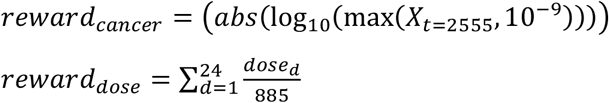

where *dose*_*d*_ is the dose belonging to the action space, body weight of all virtual patients was considered a constant (88.5 kg) and an optimal value of the constant *c* = 11 was considered.

The cancer cell count was represented in billion cells and anything below 10^-9^ was considered as 10^-9^ as it corresponds to one cell and the same was incorporated in the *reward*_*cancer*_ term. According to the defined *reward*_*cancer*_ term, a positive reward in the ranges of 5 to 9 was assigned for cancer cell count less than the desired value of 10^-5^. While the cancer cell counts greater than 10^-5^ attracts a negative reward in *reward*_*cancer*_ in the ranges of -13 to -6. The second component of the reward function, *reward*_*dose*_ is a penalty term which is defined as the normalized total drug administered to a patient.

#### Proximal Policy Optimization

Proximal policy optimization, a gradient-based reinforcement learning algorithm, was used to train the RL model. Schulman *et al*. introduced PPO ^56^ to improve training stability and restrict the policy change at each update using clipped objective function. A brief description of the algorithm is provided here, with further details available in^57^. PPO employs two neural networks: an actor and a critic network. The actor network learns the policy function *π*_*θ*_ (*S, a*), while the critic network assess the learned policy utilizing the estimated state-value function 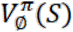 . Additionally, PPO uses a hyperparameter ϵ to prevent large policy updates by restricting the probability ratio 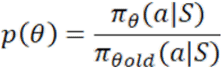 in (*1-ϵ, 1+ϵ*). In each iteration(m) of the training, L number of finite length trajectories are selected using the current policy and for each trajectory, N dosing decisions are simulated. Accordingly, the value function is generally estimated in the form of discounted reward of the trajectory as presented in equation(2).

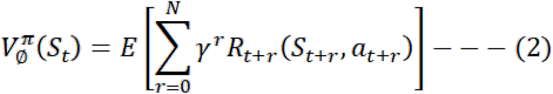

Where *R*_*t*_ is the reward received by the agent for action *a*_*t*_ at state *S*_*t*_ and transitioning to state *S*_*t*+1_ and γ ∈ [0,1] is discounting factor. As previously mentioned, in this study, a terminal reward is used instead of computing reward at each decision point *t*.

The performance of the dosing policy π_θ_ is assessed through the advantage function (equation (3)), which provides feedback to the policy network and drives gradient-based updates of the policy parameters toward optimality. Generalized advantage estimation technique is employed to compute the advantage function associated with a policy truncated to T time steps^58^.

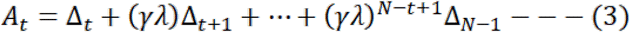

Where Δ_*t*_ *= R*_*t*_ + *γV(S*_*t*+1_ ) − *V*(*S*_*t*_ ) and λ ∈ [0,1] is used as a weighting parameter.

Finally, the parameters of actor and critic networks, θ and *ϕ*, are updated separately as per equation (4) and (5):

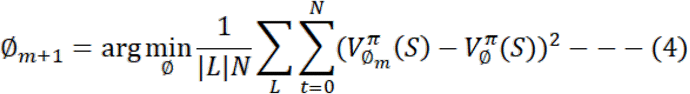

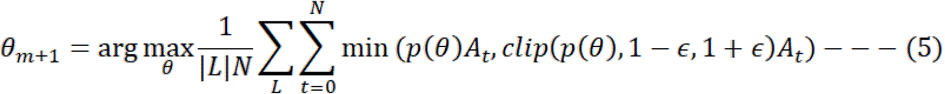

### Monte Carlo Tree Search

Monte Carlo tree search is a sampling-based planning algorithm designed to support decision-making in large, complex search spaces. Rather than exhaustively exploring all possible action sequences, MCTS incrementally constructs a partial search tree by repeatedly simulating trajectories from the current state. Each MCTS iteration follows a cycle consisting of node selection, tree expansion, rollout, and backpropagation of reward. Action selection during tree traversal is typically guided by a bandit-inspired criterion that balances exploitation of high-value actions with exploration of insufficiently sampled alternatives^53,59^. MCTS is integrated in the proposed dose optimization framework to further improve the generalized policy learned by the RL model by enabling patient specific refinement through simulation of the consequences of the various actions for multiple future decision points. To perform this task, MCTS generates a search tree in which the state St is considered as node and action at is considered as edge^53^. Each child node of St are the states resulting from applying the possible actions on St. MCTS iteratively expands the asymmetric search tree, with exploration guided by an action-selection policy that is updated iteratively to favor high-potential directions^59^. For a given node corresponding to a state the search is initialized by considering a set of five candidate actions, comprising two randomly sampled actions and three actions suggested by the trained RL policy. Each candidate is evaluated via rollout simulations over a fixed horizon of one year, followed by backpropagation of the accumulated reward to update node values. The optimal action is then selected based on these evaluations. Considering the different goal of MCTS and RL, a modified reward function is used for MCTS and is provided in equation (6).

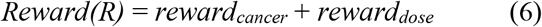

if (*X*_*t+365*_ > 10^-5^):

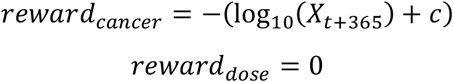

else:

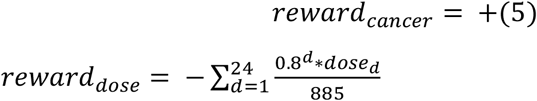

### Implementation Details

The PPO algorithm was implemented using the Stable-Baselines3 MlpPolicy, corresponding to a PyTorch-based actor–critic multilayer perceptron embedded within a custom Gym-compatible QSP environment. For the vector-based QSP state input, the PPO policy used the standard Stable-Baselines3 actor–critic network, with separate actor and critic network after input flattening. Each network comprised two fully connected hidden layers with 64 neurons per layer and tanh nonlinearities^60^. The output layer of the actor network parameterized a categorical distribution over the 11 discrete dose actions, whereas the output layer of the critic network produced a scalar state-value function. Model training was performed using the Adam optimization algorithm with mini-batch stochastic gradient descent having batch size of 128, learning rate of 3*10^-4^, discount factor of 0.99, entropy coefficient of 0.1 and 750,000 training time steps. The complete computational framework was implemented in Python except for the QSP model which was implemented in Julia. The PPO-based RL model was implemented using the PyTorch-based Stable-Baselines3 library, whereas MCTS was implemented as a Python-based inference-time planning module coupled to the trained PPO policy.

## Notes

### Competing Interest Statement

The authors have declared no competing interest.

