## SupplementaryFile for "Mechanistically informed adaptive dosing for cancer immunotherapy using AI-guided decision making"

[1]TCS Research (Life Sciences division), Tata Consultancy Services, Hyderabad, 500081, India.

**Table T1: Kolmogorov-Smirnov statistic for the cell types in different virtual population generation methods compared with the reference population.**

| | LHS | | LHS-N | | LHS-MN | | Allen's method | |
|---|---|---|---|---|---|---|---|---|
| | KS Stats | p-val | KS Stats | p-val | KS Stats | p-val | KS Stats | p-val |
| $T_{cyto}$ | 0.62 | 2.9E-87 | 0.24 | 4.3E-12 | 0.23 | 1.3E-11 | 0.22 | 5.7E-10 |
| $T_{reg}$ | 0.33 | 2.1E-23 | 0.04 | 0.80 | 0.07 | 0.240 | 0.02 | 1.0E+00 |
| **Dendritic** | 0.45 | 5.2E-44 | 0.08 | 0.09 | 0.12 | 0.003 | 0.06 | 3.4E-01 |
| **Cancer** | 0.26 | 6.8E-15 | 0.06 | 0.39 | 0.05 | 0.495 | 0.06 | 3.6E-01 |
| $T_{cyto}$**:Dendritic** | 0.53 | 1.0E-61 | 0.22 | 1.3E-10 | 0.11 | 0.005 | 0.09 | 4.6E-02 |
| $T_{cyto}$:$T_{reg}$ | 0.52 | 1.9E-60 | 0.23 | 1.3E-11 | 0.22 | 3.5E-10 | 0.22 | 6.1E-10 |
| $T_{reg}$**:_Dendritic** | 0.17 | 1.9E-06 | 0.06 | 0.31 | 0.22 | 1.2E-10 | 0.25 | 2.4E-12 |
| **Cancer:**$T_{cyto}$ | 0.44 | 3.8E-43 | 0.17 | 1.0E-06 | 0.04 | 0.88 | 0.09 | 7.0E-02 |
| **Cancer:**$T_{reg}$ | 0.05 | 0.696 | 0.09 | 0.04 | 0.17 | 1.4E-06 | 0.17 | 2.2E-06 |
| **Cancer:Dendritic** | 0.08 | 0.068 | 0.06 | 0.26 | 0.05 | 0.52 | 0.08 | 1.4E-01 |

**Table T2: Pearson's correlation between the features of the virtual patients for all the virtual population generation methods are compared with the reference population.**

| LHS | | | | | |
|---|---|---|---|---|---|
| | $T_{cyto}$ | $T_{reg}$ | **Dendritic** | **Cancer** | **IL-2** |
| $T_{cyto}$ | 1.00 | 0.02 | 0.02 | 0.01 | -0.04 |
| $T_{reg}$ | 0.02 | 1.00 | -0.02 | -0.01 | -0.02 |
| **Dendritic** | 0.02 | -0.02 | 1.00 | 0.05 | 0.01 |
| **Cancer** | 0.01 | -0.01 | 0.05 | 1.00 | 0.01 |
| **IL-2** | -0.04 | -0.02 | 0.01 | 0.01 | 1.00 |
| **LHS-N** | | | | | |
| | $T_{cyto}$ | $T_{reg}$ | **Dendritic** | **Cancer** | **IL-2** |
| $T_{cyto}$ | 1.00 | 0.07 | 0.01 | 0.00 | 0.01 |
| $T_{reg}$ | 0.07 | 1.00 | -0.01 | -0.04 | 0.02 |
| **Dendritic** | 0.01 | -0.01 | 1.00 | -0.05 | 0.00 |
| **Cancer** | 0.00 | -0.04 | -0.05 | 1.00 | -0.01 |
| **IL-2** | 0.01 | 0.02 | 0.00 | -0.01 | 1.00 |

| LHS-MN | | | | | |
|---|---|---|---|---|---|
| | $T_{cyto}$ | $T_{reg}$ | Dendritic | Cancer | IL-2 |
| $T_{cyto}$ | 1.00 | 0.02 | 0.18 | -0.11 | -0.01 |
| $T_{reg}$ | 0.02 | 1.00 | 0.21 | -0.64 | 0.04 |
| Dendritic | 0.18 | 0.21 | 1.00 | -0.52 | 0.09 |
| Cancer | -0.11 | -0.64 | -0.52 | 1.00 | -0.07 |
| IL-2 | -0.01 | 0.04 | 0.09 | -0.07 | 1.00 |
| Allen's Method | | | | | |
| | $T_{cyto}$ | $T_{reg}$ | Dendritic | Cancer | IL-2 |
| $T_{cyto}$ | 1.00 | 0.02 | 0.01 | 0.04 | 0.00 |
| $T_{reg}$ | 0.02 | 1.00 | -0.03 | 0.00 | 0.05 |
| Dendritic | 0.01 | -0.03 | 1.00 | 0.07 | -0.04 |
| Cancer | 0.04 | 0.00 | 0.07 | 1.00 | -0.02 |
| IL-2 | 0.00 | 0.05 | -0.04 | -0.02 | 1.00 |

**Table T3: Covariance between the features of the virtual patients for all the virtual population generation methods are compared with the reference population.**

| LHS | | | | | |
|---|---|---|---|---|---|
| | $T_{cyto}$ | $T_{reg}$ | Dendritic | Cancer | IL-2 |
| $T_{cyto}$ | 0.001371 | 4.7E-05 | 9E-05 | 2.5E-05 | 1.2E-05 |
| $T_{reg}$ | 4.7E-05 | 0.005337 | -0.000183 | -9.7E-05 | -2.3E-05 |
| Dendritic | 9E-05 | -0.000183 | 0.012978 | 0.000709 | -0.000511 |
| Cancer | 2.5E-05 | -9.7E-05 | 0.000709 | 0.016925 | 1.1E-05 |
| IL-2 | 1.2E-05 | -2.3E-05 | -0.000511 | 1.1E-05 | 0.015487 |
| LHS-N | | | | | |
| | $T_{cyto}$ | $T_{reg}$ | Dendritic | Cancer | IL-2 |
| $T_{cyto}$ | 0.000185 | 0.000038 | 0.000005 | -0.000002 | 0.000002 |
| $T_{reg}$ | 0.000038 | 0.001708 | -0.000020 | -0.000150 | -0.000012 |
| Dendritic | 0.000005 | -0.000020 | 0.002406 | -0.000218 | 0.000321 |
| Cancer | -0.000002 | -0.000150 | -0.000218 | 0.006597 | 0.000895 |
| IL-2 | 0.000002 | -0.000012 | 0.000321 | 0.000895 | 0.003532 |
| LHS-MN | | | | | |
| | $T_{cyto}$ | $T_{reg}$ | Dendritic | Cancer | IL-2 |
| $T_{cyto}$ | 0.000180 | 0.000011 | 0.000115 | -0.000116 | -0.000016 |
| $T_{reg}$ | 0.000011 | 0.001614 | 0.000409 | -0.002014 | 0.000231 |
| Dendritic | 0.000115 | 0.000409 | 0.002366 | -0.001983 | 0.000620 |
| Cancer | -0.000116 | -0.002014 | -0.001983 | 0.006232 | -0.000737 |
| IL-2 | -0.000016 | 0.000231 | 0.000620 | -0.000737 | 0.020186 |
| Allen's Method | | | | | |
| | $T_{cyto}$ | $T_{reg}$ | Dendritic | Cancer | IL-2 |

| | | | | | |
|---|---|---|---|---|---|
| **$T_{cyto}$** | 0.000196 | 0.000010 | 0.000004 | 0.000049 | 0.000001 |
| **$T_{reg}$** | 0.000010 | 0.001860 | -0.000063 | 0.000010 | 0.000281 |
| **Dendritic** | 0.000004 | -0.000063 | 0.002500 | 0.000317 | -0.000307 |
| **Cancer** | 0.000049 | 0.000010 | 0.000317 | 0.007627 | -0.000234 |
| **IL-2** | 0.000001 | 0.000281 | -0.000307 | -0.000234 | 0.020130 |

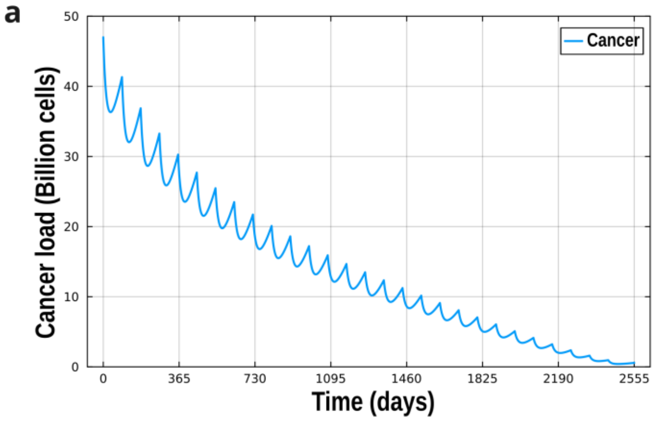

D1 = [3, 3, 3, 3, 3, 3, 3, 3, 3, 3, 3, 3, 3, 3, 3, 3, 3, 3, 3, 3, 3, 3, 3, 3, 3, 3, 3, 3]
Total drug = 84 mg/kg

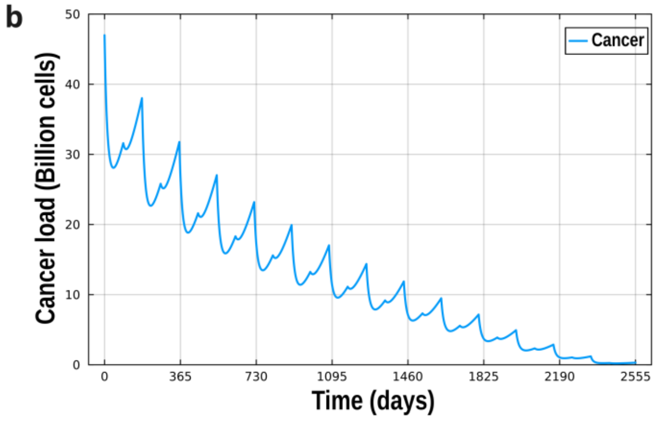

D2 = [5, 1, 5, 1, 5, 1, 5, 1, 5, 1, 5, 1, 5, 1, 5, 1, 5, 1, 5, 1, 5, 1, 5, 1, 5, 1, 5, 1]
Total drug = 84 mg/kg

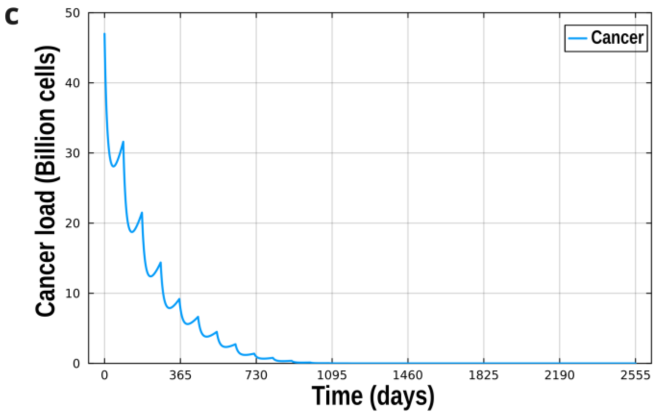

D3 = [5, 5, 5, 5, 4, 4, 4, 4, 3, 3, 3, 3, 3, 3, 3, 3, 3, 3, 3, 3, 2, 2, 2, 2, 1, 1, 1, 1]
Total drug = 84 mg/kg

**Figure S1: The effect of dosing pattern on the cancer progression outcome is shown for an example virtual patient for three different dosing regimens. The total drug administered in all three cases is same (84 mg/kg) while the pattern of dosing is different. The initial state of the patient, parameters, drug (*ipilimumab*) and dosing interval (180 days) are consistent in all three simulations. The progression of cancer load over time significantly changes as the dosing pattern is changed. (a) D1: a constant dose of 3 mg/kg is administered in all the 28 dosing time points (b) D2: Alternate doses of a high dose of 5 mg/kg followed by by a low dose of 1 mg/kg is administered (c) D3: Starting with high dose of 5mg/kg, the dose is gradually reduced in the subsequent dosing timepoints.**

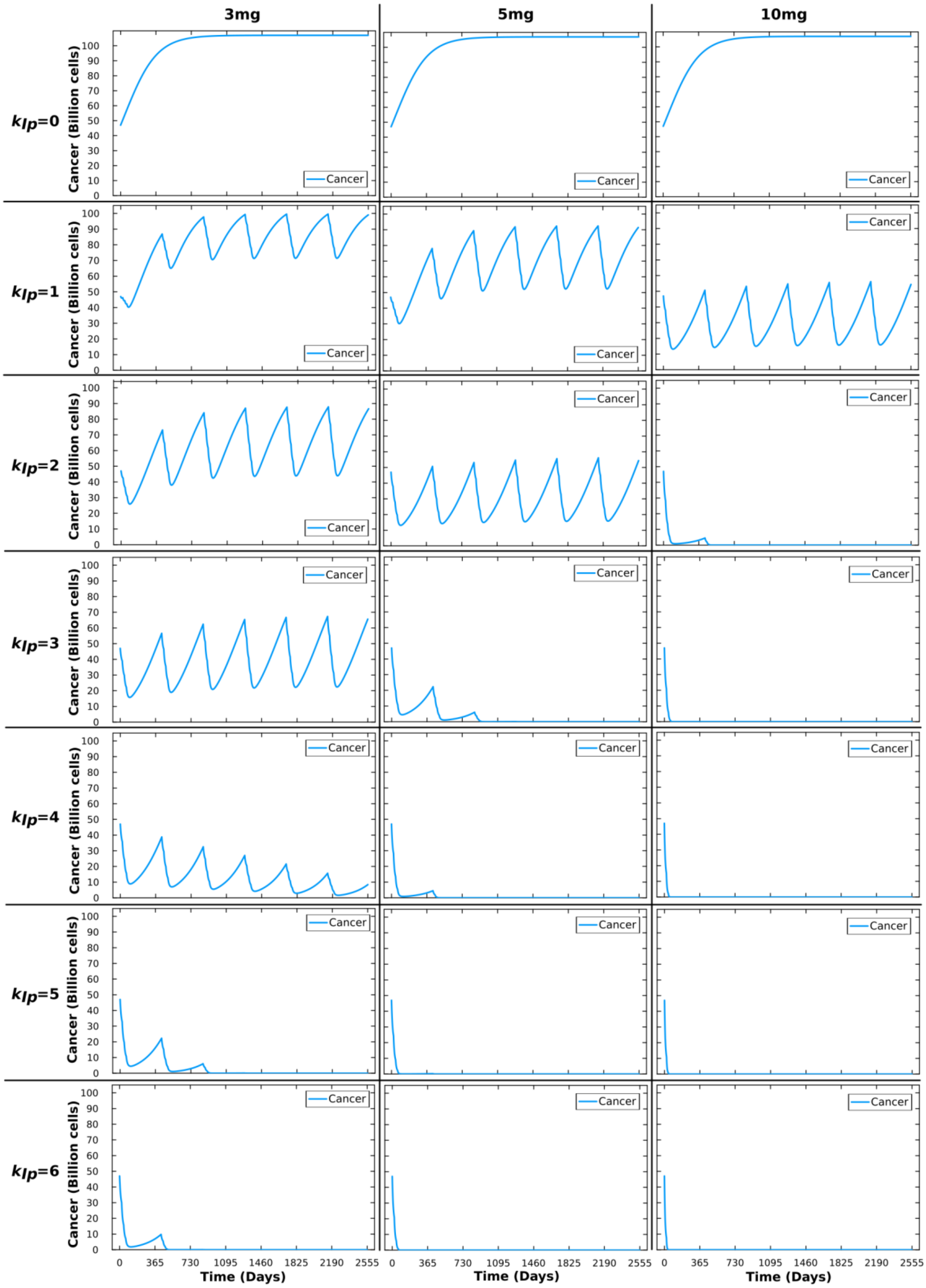

**Figure S2: The cancer progression is simulated for 7 years by changing only the values of the parameter $k_{Ip}$ (killing rate of cancer cells by CTL due to *ipilimumab*) for three different fixed dosing of the drug *ipilimumab* (3mg/kg, 5mg/kg and 10mg/kg). All the parameters and initial state variables (cell counts and molecular concentrations) are kept constant. The experiment shows the effect of $k_{Ip}$ on the outcome of the drug administration. $k_{ip}$ value of 0 means that there is no effect of the drug and cancer cells are not killed**

**by cytotoxic T-cells due to the drug *ipilimumab*. In that case, the cancer load increases to the highest saturation state in all three dosing protocols. As $k_{ip}$ increases, the effect of the drug can be observed. Remission is observed for 10mg/kg dose when $k_{ip}$ value of 2, whereas for 5mg/kg and 3mg/kg the remission is attained at $k_{ip}$ values of 3 and 5 respectively. Generating virtual patients with $k_{ip}$ values in the range of 0 to 6 ensures coverage of the complete response profiles in the virtual population.**

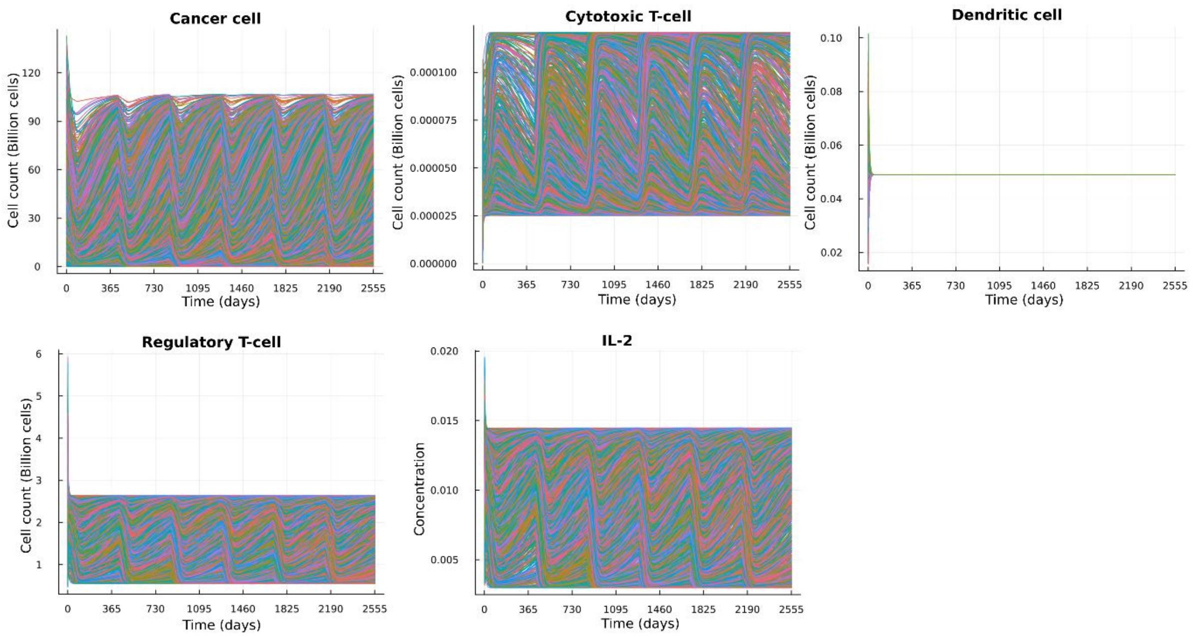

**Figure S3: The temporal profile of the four cell types and signalling molecule generated by QSP simulation of the 10,000 virtual patients (VPop) for a period of 7 years. The simulation was carried out by administering a constant dose of 3mg/kg body weight for yearly 4 doses at an interval of 3 weeks followed by a gap of one year. The variables remain within the bounds for all the patients which shows that the generated virtual population is stable. The four cell types considered here include cancer cells, cytotoxic T-cells, dendritic cells and regulatory T-cells, and the signalling molecule considered is IL-2.**

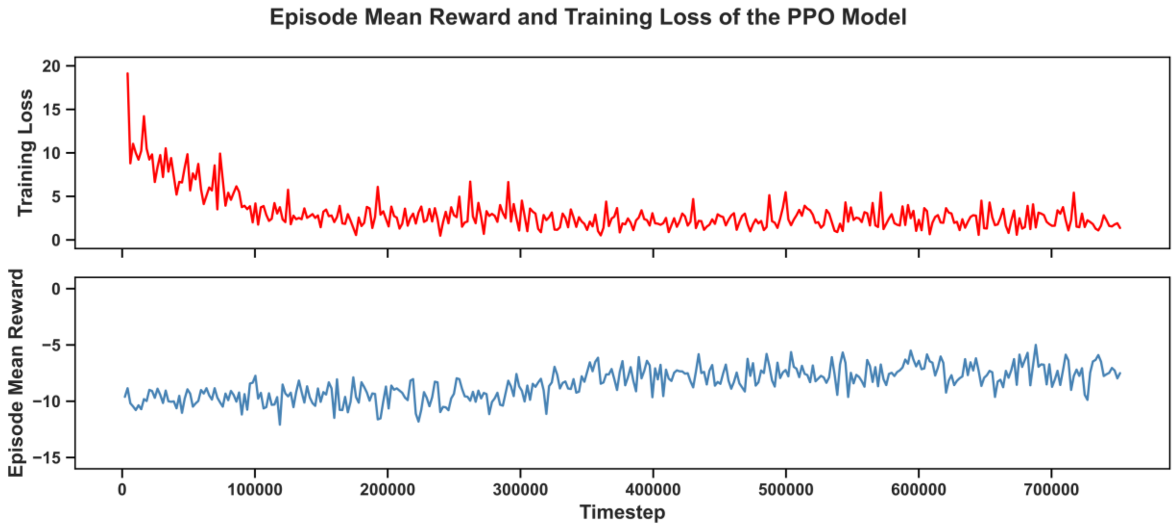

**Figure S4: Plot of training loss and episode mean reward over during the RL model training.**

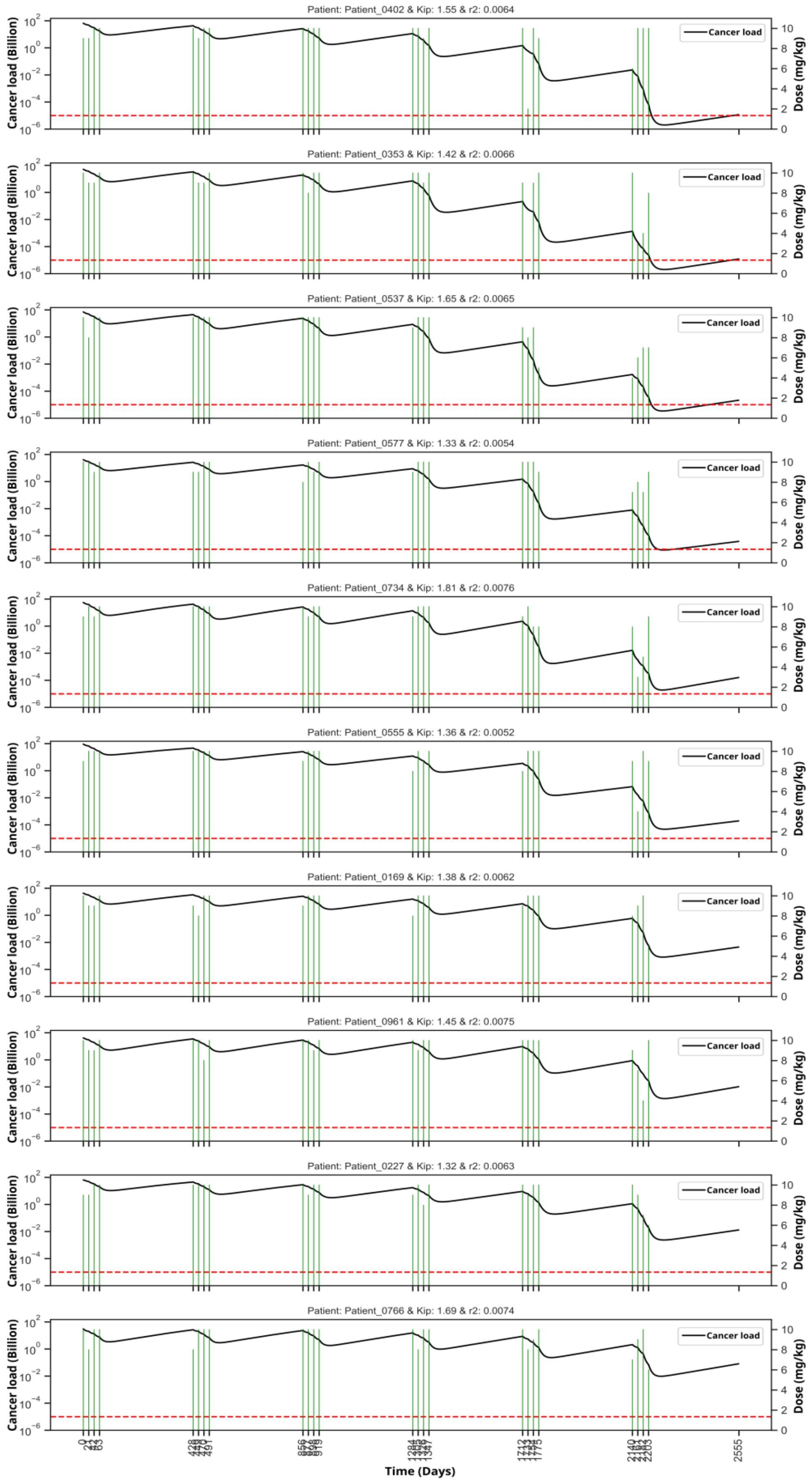

**Figure S5: Plot of cancer progression and dose administration for 10 patients that were not remitted by the final model but were remitted by the 10 mg/kg fixed dosing regimen.**

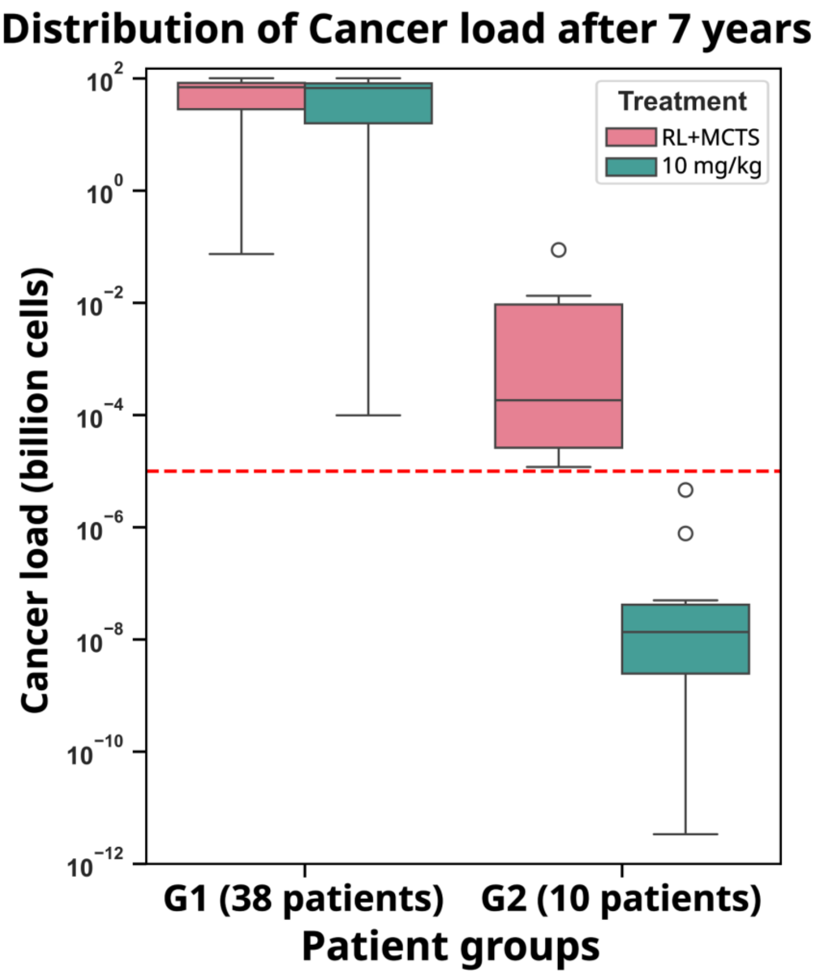

**Figure S6: Distribution of cancer load at the end of 7 years is shown for virtual patients in the test set (*$VPop_{Test}$*) that were not remitted by the final model. These 48 patients are divided into two groups – G1 of 38 patients that were not remitted even by the 10 mg/kg dosing regimen and G2 of 10 patients that were remitted by the 10 mg/kg regimen.**
